# A convergent metabolic-kinase signaling axis links Parkinson’s disease and multiple system atrophy

**DOI:** 10.64898/2026.06.11.731777

**Authors:** Erika Velasquez, Ekaterina Savchenko, Alejandro Gomez Toledo, Elin Näsström, Annika I. Johansson, Arianna Colini Baldeschi, Tomas Deierborg, Katie Lunnon, Yuriy Pomeshchik, Melinda Rezeli, Laurent Roybon

## Abstract

Parkinson’s disease (PD) and multiple system atrophy (MSA) are alpha-synucleinopathies with overlapping clinical phenotypes but distinct cellular pathological hallmarks. Whether these disorders share upstream molecular changes beyond alpha-synuclein aggregation remains unresolved. Here, we generated induced pluripotent stem cell (iPSC)-derived midbrain spheroids containing dopaminergic neurons from individuals with monogenic PD, idiopathic PD, MSA, and controls, and applied integrated proteomics, metabolomics, and phosphoproteomics to define disease-associated biochemical programs and their regulatory architecture. Despite their distinct etiologies, PD and MSA spheroids displayed highly concordant molecular remodeling, with cellular metabolism emerging as the dominant shared disturbance. Network analyses identified coordinated changes in central carbon metabolism, oxidative phosphorylation, branched-chain amino acid catabolism, pantothenate/CoA metabolism, and lipid remodeling, coupled to phosphorylation-driven rewiring of MAPK, mTOR, AMPK, PKA/PKC, and second-messenger kinase programs. Importantly, these metabolic and signaling axes were also prominent in postmortem substantia nigra from PD and MSA donors, supporting conservation between patient-derived models and postmortem brain tissue. Together, these data identify metabolic dysregulation as a unifying molecular feature across PD and MSA and suggesting phosphorylation-linked metabolic control nodes as candidate entry points for therapeutic intervention.

**Highlights:**

1. PD and MSA exhibit molecular programs that extend beyond alpha-synuclein pathology.
2. Proteomics identifies metabolism as the dominant shared disease axis across PD and MSA.
3. Metabolomics resolves a coupled glucose-TCA-BCAA-CoA-lipid remodeling program in PD and MSA.
4. Phosphorylation-centered kinase networks link signaling rewiring to metabolic bottlenecks.
5. Shared metabolic features are conserved between patient iPSC-derived midbrain spheroids and substantia nigra.

**Graphic abstract (generated using BioRender):** 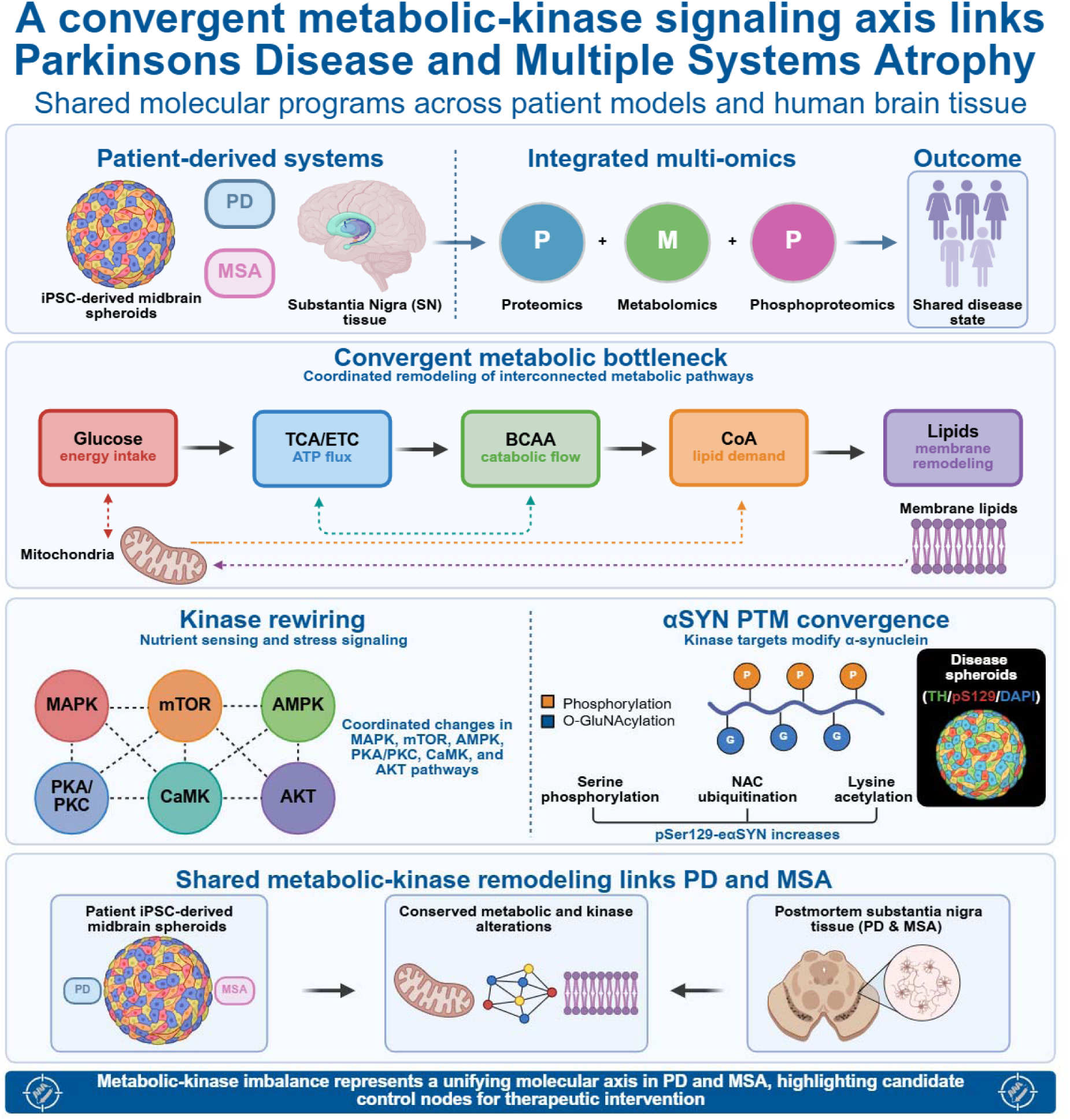

## 1. Introduction

Parkinson’s disease (PD) and multiple system atrophy (MSA) are progressive neurodegenerative disorders classified as synucleinopathies, defined by misfolded alpha-synuclein (aSYN) accumulation accompanied by dopamine neuron degeneration^1,2^. PD pathology is characterized by intraneuronal Lewy bodies and Lewy neurites, whereas MSA is classically defined by aSYN-positive oligodendroglial cytoplasmic inclusions^3^. Despite this divergence in cellular pathology, PD and MSA share overlapping motor and non-motor phenotypes and can be difficult to distinguish clinically, particularly at early stages^4,5^. These observations raise the possibility that PD and MSA converge on upstream molecular drivers beyond protein aggregation, but direct molecular comparisons remain limited.

Multiple cellular mechanisms have been implicated in PD and/or MSA pathogenesis, including increased SNCA expression^6,7^, aggregation-promoting post-translational modifications^8^, disease-associated genetic variants^9,10^, and impaired aSYN clearance pathways^11^. Beyond aSYN-centered mechanisms, prior studies have reported dysfunction across several cellular systems, including altered autophagy-lysosomal function^12–14^, disrupted ubiquitin-proteasome system activity^15–17^, mitochondrial dysfunction and impaired bioenergetics^18–20^, membrane trafficking perturbations^21^, and inflammatory signaling^22,23^. However, most molecular studies have examined PD and MSA separately or focused on a single molecular layer, such as the transcriptome, proteome, or metabolome^24–29^. As a result, it remains unclear which biochemical programs are shared between PD and MSA, which are disease-specific, and how upstream regulatory networks shape downstream metabolic states.

Integrating protein abundance with metabolite readouts and phosphorylation-dependent signaling is essential for resolving the regulatory architecture of complex neurodegenerative processes. Human induced pluripotent stem cell (iPSC)–based models offer a controlled framework for examining early, cell-intrinsic disease-associated remodeling while minimizing environmental confounders^30^. However, linking model-derived molecular states to human disease requires direct comparison with postmortem brain tissue, where end-stage pathology and cellular composition can amplify disease-specific divergence.

Here, we applied a systems-level multi-omics strategy across iPSC-derived midbrain spheroids containing dopaminergic neurons and postmortem substantia nigra from PD, MSA, and control donors. We identify striking convergence between PD and MSA on a core metabolic axis spanning central carbon metabolism, branched-chain amino acid (BCAA)/CoA handling, and lipid remodeling, and delineate phosphorylation-dependent kinase networks that map onto these metabolic bottlenecks. Importantly, the found the same metabolic signatures prominent in human substantia nigra, supporting convergence between patient-derived models and human disease state.

## 2. Results

### 2.1. PD and MSA iPSC-derived midbrain spheroids show comparable dopaminergic differentiation and a shared disease-associated proteomic signature

To enable direct molecular comparison of PD and MSA across etiologies, we generated iPSC-derived midbrain spheroids from familial PD lines carrying GBA variants (n = 3), LRRK2 variants (n = 3), PARK2 variants (n = 3), PINK1 variants (n = 3), and SNCA multiplications (n = 4), as well as from individuals with idiopathic PD (n = 3), MSA (n = 3; MSA-P in this cohort), and controls (n = 3) (Table S1). Spheroids were differentiated using a staged ventral midbrain patterning and maturation protocol and aged to day 100 for downstream profiling (Figure 1A). All samples were sequentially examined using proteomics, metabolomics, and phosphoproteomics, and the resulting datasets were integrated through a harmonized multi-layer analytical workflow incorporating normalization, covariate-aware batch correction, pathway enrichment, and network analyses (Figure S1).

**Figure 1.**
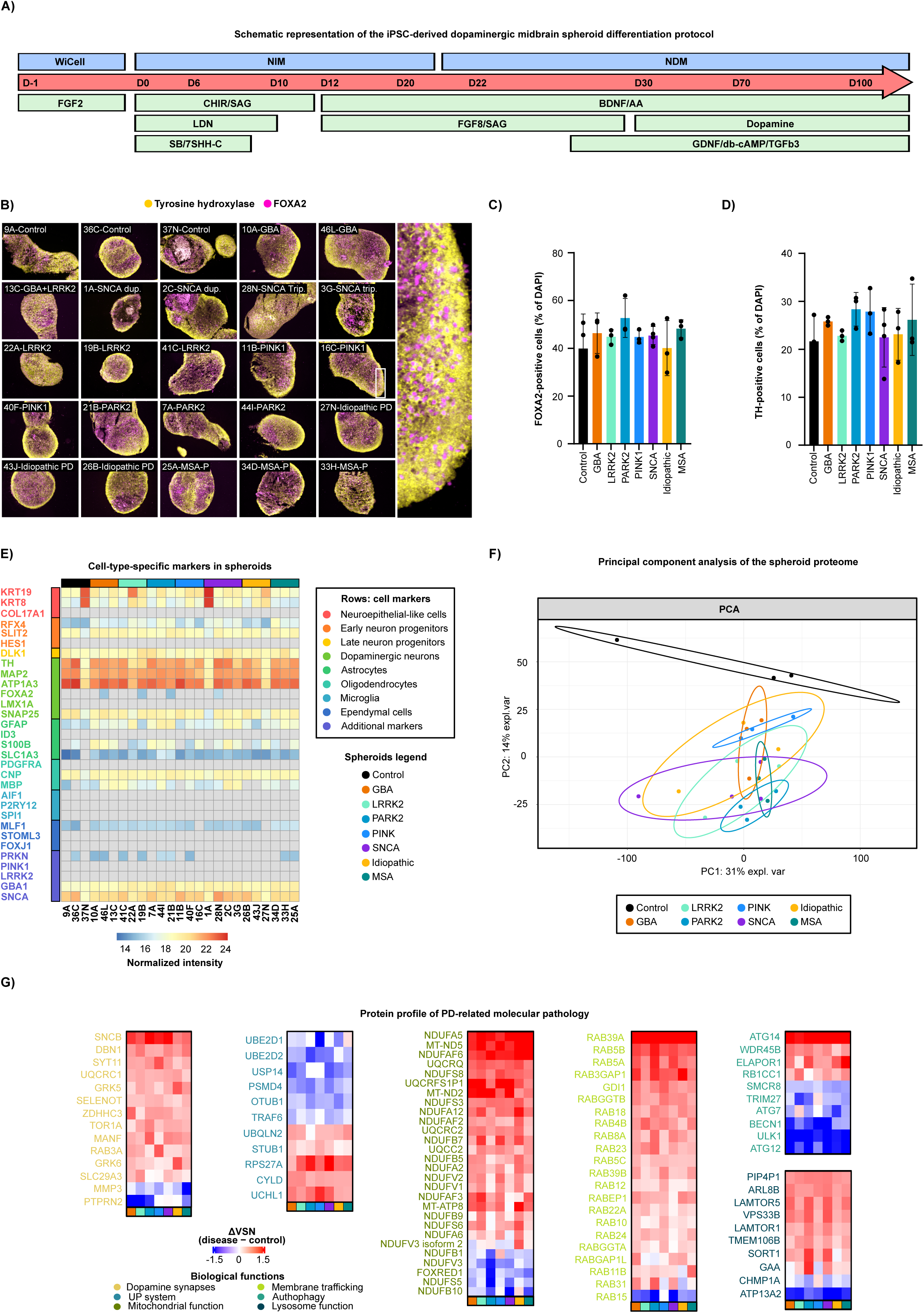
Generation and proteomic characterization of iPSC-derived dopaminergic midbrain spheroids in PD and MSA. (A) Schematic representation of the iPSC-derived dopaminergic midbrain spheroid differentiation protocol. iPSC lines from GBA (N=3; 2♂ and 1♀), LRRK2 (N=3; 2♂, 1♀), PARK2 (N=3; 0♂, 3♀), PINK (N=3; 2♂, 1♀), SNCA (N=4; 0♂, 4♀), Idiopathic PD (N=3; 2♂ and 1♀), MSA (N=3; 1♂, 2♀) and controls (N=3; 2♂, 1♀) were differentiated into midbrain spheroids containing dopaminergic neurons. Spheroids were collected at day 100, corresponding to mature/aged dopaminergic neuronal and glial populations, and processed for downstream analyses. (B) Immunostaining for the midbrain marker FOXA2 and the dopaminergic neuron marker tyrosine hydroxylase. (C–D) Bar graphs showing the percentage of FOXA2-positive(C) and TH-positive cells (D) among DAPI-positive cells. (E) Assessment of cell-type-specific markers in spheroids based on normalized protein intensities derived from the proteomics dataset. No significant differences were observed across groups. (F) Principal component analysis (PCA) of the spheroid proteomes. The PCA plot shows clear separation between control and disease conditions, whereas substantial overlap is observed between PD and MSA spheroids. (G) Annotated heatmaps of PD-associated proteomic alterations. Differentially expressed proteins relative to controls (robust empirical Bayes, p < 0.05, |log2FC| > ±0.5) are organized according to biological pathways previously implicated in PD pathology.

Immunostaining at day 100 confirmed robust acquisition of ventral midbrain identity and dopaminergic differentiation across all genotypes and diagnoses, with widespread expression of FOXA2 and tyrosine hydroxylase (TH; Figure 1B). The proportions of FOXA2^+^ and TH^+^ cells did not reveal systematic differences among groups (Figure 1C and D), indicating comparable dopaminergic differentiation across lines. To evaluate cellular composition beyond histology, we leveraged proteome-based marker profiling. A curated panel^24^ spanning neuroepithelial-like markers (e.g., KRT8/KRT19, COL17A1), neuronal progenitor markers (e.g., RFX4, HES1, DLK1), dopaminergic and neuronal proteins (e.g., TH, LMX1A, MAP2, SNAP25, ATP1A3, FOXA2), and glial-lineage markers (e.g., GFAP, S100B, SLC1A3, CNP, MBP) showed highly similar marker abundance profiles across spheroids. Microglia-associated proteins (e.g., AIF1, P2RY12, SPI1) were included in the panel and were largely absent from the spheroid proteome, consistent with limited microglial contribution under this differentiation paradigm (Figure 1E; Table S2). Together with the immunostaining data, these analyses support comparable marker-defined cellular composition across PD, MSA, and control spheroids at day 100.

In addition to a similar cellular composition, global proteomics (Table S3) revealed a shared disease-associated state. Principal component analysis (PCA) of the spheroid proteome separated controls from patient-derived spheroids, with PD and MSA clustering closely together (PC1: 31% explained variance; PC2: 14%; Figure 1F). We further observed dysregulation of protein groups (Table S4) previously linked to PD-relevant biology, including mitochondrial respiratory chain and assembly factors^18^(e.g., NDUF and related proteins), ubiquitin-proteasome components^31^ (e.g., USP14, PSMD4, UBE2D family), membrane/vesicle trafficking regulators^32^ (extensive RAB network proteins), and autophagy/lysosomal factors^33^ (e.g., ULK1, ATG14, lysosome-associated proteins; Figure 1G). Dopamine synapse-associated proteins (e.g., SNCB^34^, SYT11^35^) were also part of the dysregulated signature, consistent with a disease-relevant dopaminergic molecular phenotype (Figure 1G). Collectively, PD and MSA iPSC-derived midbrain spheroids exhibit reproducible dopaminergic differentiation with broadly similar marker-defined composition, yet share a robust disease-associated proteomic signature suitable for mechanistic analysis of convergent synucleinopathy biology.

### 2.2. Metabolic remodeling emerges as the dominant shared program across PD and MSA spheroids

To define the biological programs underlying the shared PD/MSA proteome, we performed pathway enrichment analysis of dysregulated proteins (Table S4) across PD subtypes and MSA. Strikingly, PD and MSA showed highly similar enriched pathways, with metabolic processes representing the most robust and recurrent enrichments. These included amino acid metabolism, carbohydrate and glucose metabolism including glycolysis and glucose regulation, lipid metabolism including phospholipid, glycerolipid, sphingolipid, and steroid-related pathways, and mitochondrial energy metabolism including TCA/ETC-linked biology (Figure 2A). Beyond metabolism, shared dysregulated programs included autophagy/mitophagy and apoptosis, ubiquitin-mediated proteolysis and protein ubiquitination, mitochondrial protein degradation, ER protein processing, ER-to-Golgi transport, Asn N-linked glycosylation, and broader membrane trafficking programs (Figure 2A). Shared signaling-associated pathways included mTOR/MAPK signaling, receptor tyrosine kinase signaling, Rho GTPase signaling, DAG/IP_3_ signaling, and immune-associated modules, including interferon- and MHC-related pathways (Figure 2A).

**Figure 2.**
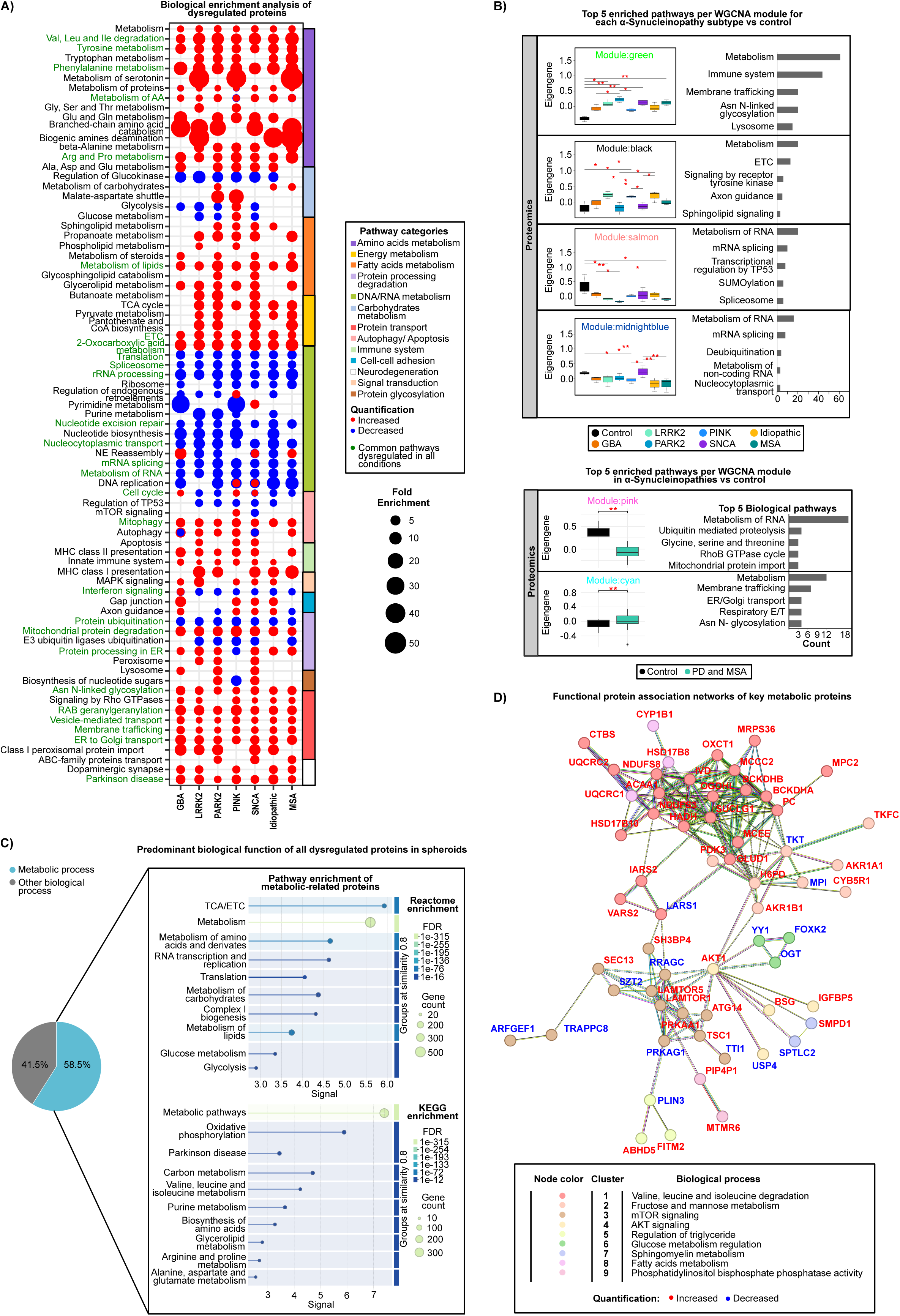
Proteomic pathway and network analyses reveal convergent metabolic dysregulation in PD and MSA spheroids. (A) Bubble plot summarizing pathway enrichment analysis (Fisher’s Exact test, p < 0.05) of proteins dysregulated in PD and MSA spheroids relative to controls (robust empirical Bayes, p < 0.05, |log2FC| > ±0.5). Pathways highlighted in green indicate those commonly dysregulated across all conditions. (B) Weighted gene co-expression network analysis (WGCNA) identifying protein modules significantly altered across PD, MSA, and control groups (Kruskal-Wallis, p<0.05; Mann–Whitney U test, p < 0.05). The top five enriched biological pathways for each major module are shown (Fisher’s exact test, p < 0.05). (C) A total of 58.5% of dysregulated proteins in PD and MSA spheroids are involved in metabolic processes. Pathway enrichment analysis was performed to further characterize metabolic alterations. (D) Functional network analysis highlighting key proteins driving metabolic dysregulation in PD and MSA spheroids.

To identify coordinated protein programs, we performed weighted gene co-expression network analysis (WGCNA). Subtype-resolved WGCNA identified modules with consistent behavior across alpha-synucleinopathy spheroids (Figure 2B; Table S4). A green module displayed coordinated upregulation across disease conditions and was enriched for metabolism, immune system pathways, membrane trafficking, Asn N-linked glycosylation, and lysosomal pathways. A black module showed elevated co-expression, particularly in LRRK2, PARK2, and idiopathic PD spheroids, and was enriched for metabolic pathways and electron transport chain programs, as well as signaling terms such as receptor tyrosine kinase signaling, axon guidance, and sphingolipid signaling. In contrast, salmon and midnightblue modules displayed coordinated downregulation across PD and MSA and were enriched for RNA metabolism, mRNA splicing, nucleocytoplasmic transport, SUMOylation, transcriptional regulation by TP53, and deubiquitination (Figure 2B). A complementary WGCNA grouping of PD + MSA versus control further resolved modules that capture shared disease programs, including a downregulated module enriched for ubiquitin-mediated proteolysis and amino acid metabolism, including glycine/serine/threonine, and another upregulated module enriched for membrane trafficking, ER/Golgi transport, respiratory electron transport, and Asn N-linked glycosylation (Figure 2B). Collectively, WGCNA highlights coordinated remodeling centered on metabolism, membrane trafficking, and proteostasis as a prominent shared feature of PD and MSA spheroids.

Across all dysregulated proteins, metabolic proteins accounted for 58.5% of alterations in PD/MSA spheroids (Figure 2C), confirming metabolism as the dominant shared proteomic program. Enrichment analysis of the metabolic subset highlighted central carbon and mitochondrial pathways, including oxidative phosphorylation and complex I-linked processes, as well as lipid metabolic programs and BCAA-related pathways (Figure 2C). A functional association network of key metabolic proteins identified an integrated cluster linking BCAA degradation proteins (e.g., BCKDHA/B, IVD), pyruvate/TCA-linked enzymes (e.g., MPC2, PC, OGDHL, OXCT1, SUCLG1), mitochondrial respiratory complex components (e.g., NDUFS3/S8, UQCRC1/2), and lipid-associated enzymes (e.g., HSD17B8/B10, ACAA1) (Figure 2D). This cluster was connected to nutrient- and growth-signaling regulators (AKT- and mTOR-linked nodes; including AKT1, PRKAA1/G1, LAMTOR, and RRAGC components) and glucose-regulatory proteins (e.g., OGT, FOX2, YY1) (Figure 2D). These findings suggest that metabolic remodeling is tightly coupled to phosphoregulatory signaling networks. Together, these data identify dysregulated metabolism as a central convergent feature of PD and MSA spheroids, interfacing with trafficking/proteostasis pathways and nutrient-sensing signaling networks.

### 2.3. Metabolomics defines a coupled axis of metabolic alterations in PD and MSA spheroids

Because metabolites provide a functional readout of pathway activity, we next performed complementary gas chromatography-mass spectrometry (GC-MS) and liquid chromatography-mass spectrometry (LC-MS) metabolomics on day-100 spheroids (Tables S5). Metabolomic profiling of iPSC-derived midbrain spheroids revealed widespread and heterogeneous metabolic alterations. Despite this heterogeneity, these alterations still converged on shared disease-associated programs at the pathway level. (Figure 3A; Table S6). Enrichment analysis of dysregulated metabolites consistently highlighted amino acid metabolism, including BCAA and glycine/serine/threonine pathways, central carbon metabolism (glycolysis/gluconeogenesis, pyruvate metabolism, and the pentose phosphate pathway), and lipid metabolism, including fatty acid oxidation and unsaturated fatty acid biosynthesis. Notably, the TCA cycle, pantothenate/CoA metabolism, and glutathione metabolism emerged as recurrently perturbed processes across disease conditions, pointing to a shared defect in mitochondrial carbon utilization and redox homeostasis (Figure 3A).

**Figure 3.**
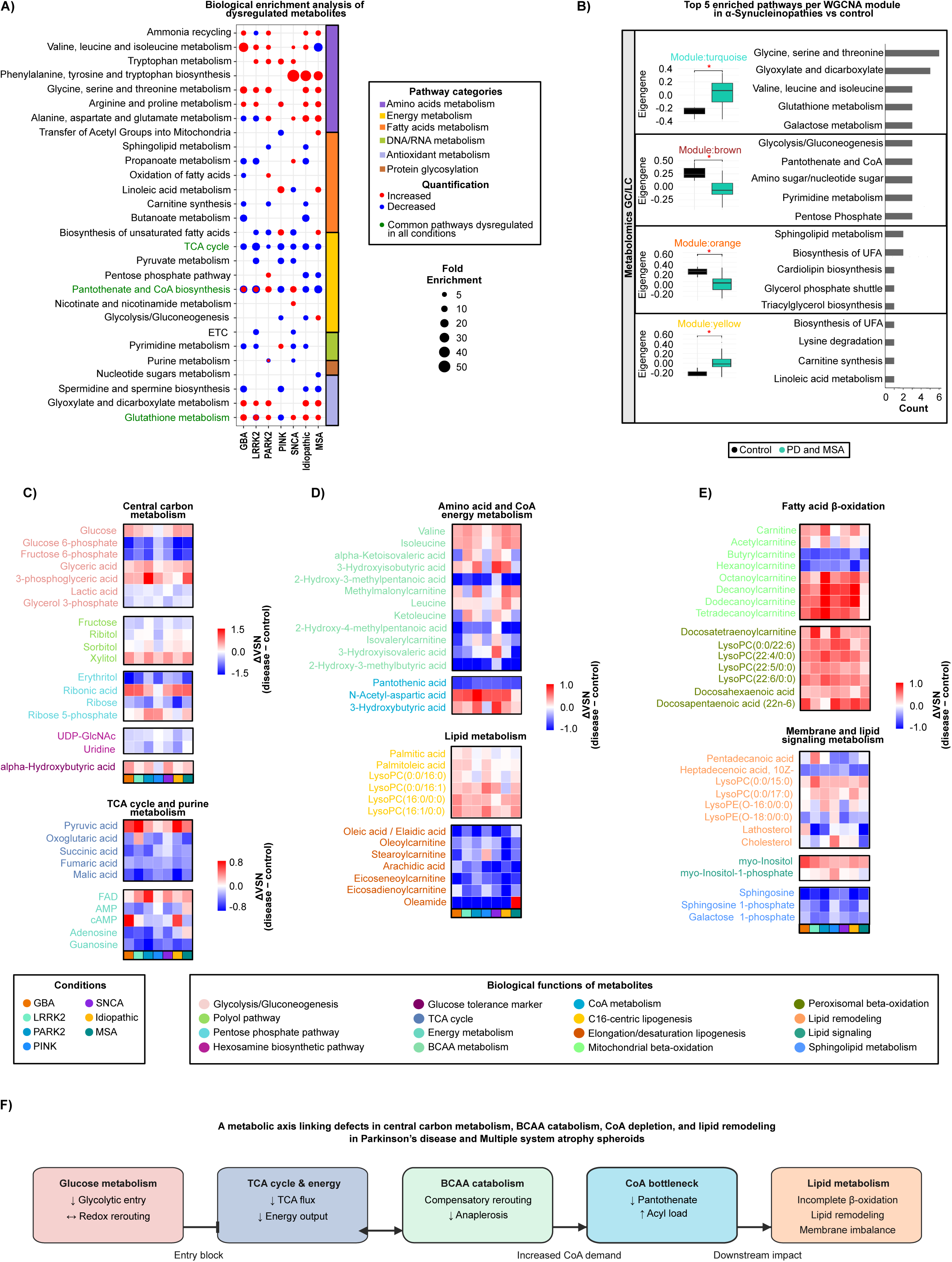
Metabolomic profiling identifies impaired carbon flux, compensatory amino acid metabolism, and lipid remodeling in PD and MSA spheroids. (A) Bubble plot summarizing pathway enrichment analysis (Fisher’s exact test, p < 0.05) of metabolites dysregulated in PD and MSA spheroids relative to controls, identified using OPLS classification (cutoff > 0.75). Pathways highlighted in green indicate those commonly dysregulated across all conditions. (B) Weighted gene co-expression network analysis (WGCNA) identifying metabolite modules significantly altered across PD, MSA, and control groups (Mann–Whitney U test, p < 0.05). The top five enriched biological pathways for each major module are shown (Fisher’s exact test, p < 0.05). (C–E) Annotated heatmaps of key dysregulated metabolites in PD and MSA spheroids, displaying disease-associated changes relative to controls (ΔVSN = disease − control). Alterations in central carbon metabolism, TCA cycle, amino acid and CoA metabolism, lipid metabolism, and membrane remodeling represent the principal metabolic axes underlying PD and MSA dysfunction. (F) Schematic model summarizing a metabolomics-defined metabolic axis in which impaired glycolytic entry restricts mitochondrial carbon flux, triggering compensatory engagement of branched-chain amino acid (BCAA) metabolism. Increased CoA demand under these conditions may create a CoA bottleneck, leading to incomplete fatty acid oxidation and consequent lipid and membrane remodeling.

To capture coordinated metabolite programs in PD plus MSA, we performed WGCNA, which identified modules linking amino acid and antioxidant metabolism with carbon routing and lipid remodeling (Figure 3B; Table S6). An upregulated turquoise module was enriched for glycine/serine/threonine metabolism, BCAA pathways, glutathione metabolism, and galactose metabolism. In contrast, a downregulated brown module captured pathways related carbon-entry and biosynthetic routing, including glycolysis/gluconeogenesis, pentose phosphate metabolism, amino sugar and nucleotide sugar metabolism, pantothenate/CoA pathways, and pyrimidine metabolism. Lipid-centered modules showed divergent directionality: a downregulated orange module linked sphingolipid metabolism, cardiolipin and triacylglycerol biosynthesis, and glycerol phosphate shuttle biology, whereas an upregulated yellow module captured carnitine synthesis, linoleic acid metabolism, and biosynthesis of unsaturated fatty acids (Figure 3B). Thus, correlation-based metabolite modules support a coordinated axis integrating carbon metabolism, amino acid routing, CoA-linked capacity, and lipid remodeling in disease spheroids.

At the metabolite level, central carbon metabolism showed coherent signatures consistent with restricted glycolytic entry and altered downstream routing. Disease spheroids accumulated glucose, whereas early glycolytic intermediates, including glucose-6-phosphate and fructose-6-phosphate, were reduced, indicating impaired entry into glycolysis (Figure 3C). Several midstream glycolytic intermediates, such as glycerate/3-phosphoglycerate and lactate-associated readouts, were also altered, consistent with disrupted glycolytic throughput (Figure 3C). In parallel, we observed changes in glucose-derived pathways, including the polyol pathway (e.g., ribitol, sorbitol, xylitol), the pentose phosphate pathway (e.g., ribose-5-phosphate, ribose, erythritol), and the hexosamine pathway (e.g., UDP-GlcNAc and uridine). These changes occurred together with increased α-hydroxybutyrate, consistent with redox-driven rerouting of glucose metabolism (Figure 3C). These observations coincided with alterations in TCA-cycle intermediates, including depletion of multiple downstream intermediates, and purine/energy-associated metabolites (including AMP-, adenosine-, guanosine-, and FAD-related features), supporting impaired mitochondrial carbon flux and perturbed energetic/redox homeostasis (Figure 3C).

A second prominent signature involved compensatory rerouting through amino-acid metabolism with impaired BCAA catabolism. Disease spheroids showed accumulation of valine, leucine, and isoleucine, together with disruption of downstream BCAA-associated intermediates, including hydroxy-acid and acylcarnitine-linked readouts, consistent with reduced oxidative flux through BCAA degradation (Figure 3D). This pattern is consistent with diminished anaplerotic support of the TCA cycle through succinyl-CoA-linked entry routes. Importantly, this BCAA-associated remodeling intersected with evidence for a CoA-limited state: disease spheroids displayed depletion of pantothenic acid (vitamin B5), the precursor for CoA biosynthesis, together with alterations in acetyl-CoA-linked proxies (e.g., N-acetyl-aspartate and 3-hydroxybutyrate patterns), consistent with a pronounced CoA bottleneck under sustained demand for acyl-group processing (Figure 3D).

Finally, glucose/BCAA disruption and CoA limitation were accompanied by broad lipid remodeling suggesting with incomplete β-oxidation and compensatory acyl handling. Disease spheroids showed accumulation of C16-centric lipid species and lysophospholipids such as LysoPC(16:0) and LysoPC(16:1)), alongside altered elongation/desaturation-associated lipids, including oleic/elaidic acid and arachidic acid-associated features (Figure 3D). We additionally observed activation of the carnitine shuttle and acyl overflow, with altered carnitine and multiple C8–C14 acylcarnitines, consistent with incomplete mitochondrial β-oxidation under conditions of limited free CoA (Figure 3E). Lipid remodeling extended to membrane-signaling metabolites, including changes in myo-inositol and myo-inositol-1-phosphate, and alterations in sphingosine/sphingosine-1-phosphate and galactose-1-phosphate, indicating extensive membrane lipid remodeling and signaling changes (Figure 3E). Together, these results define a tightly coupled metabolic program linking impaired glycolytic entry, reduced mitochondrial/TCA output, BCAA catabolic defects, pantothenate-linked CoA limitation, and lipid remodeling in PD and MSA spheroids (Figure 3F).

### 2.4. Metabolic alterations are linked to phosphorylation-driven signaling remodeling in PD and MSA spheroids

Given that phosphorylation is a rapid and reversible mechanism for regulating metabolic routing and stress adaptation, we next examined phosphorylation-driven signaling changes accompanying the metabolic disturbances identified in PD and MSA spheroids by integrating global proteomics with phosphoproteomics and kinase-substrate inference. We first quantified, by global proteomics, phosphotransferases and phosphatases linked to metabolic control and lipid homeostasis, revealing coordinated remodeling across PD and MSA spheroids (Figure 4A). Within metabolic flux control nodes, abundance changes included increased levels of key glycolytic enzymes (HK1, PFKP, PGK1) together with increased PDK1/PDK3, a pattern predicted to inhibit pyruvate dehydrogenase activity and restrict conversion of pyruvate to acetyl-CoA. This is consistent with the pyruvate and TCA cycle depletion patterns observed by metabolomics (Figure 3C). In parallel, increased branched-chain ketoacid dehydrogenase kinase (BCKDK) supports phosphorylation-mediated inhibition of BCAA degradation, aligning with BCAA accumulation and downstream catabolic disruption (Figure 3D). We also observed increased abundance of enzymes linked to CoA and lipid metabolism, including COASY, MVK, and PMVK, suggesting a compensatory attempt to sustain acetyl-CoA/CoA-linked metabolism and lipid homeostasis under metabolic stress (Figure 4A).

**Figure 4.**
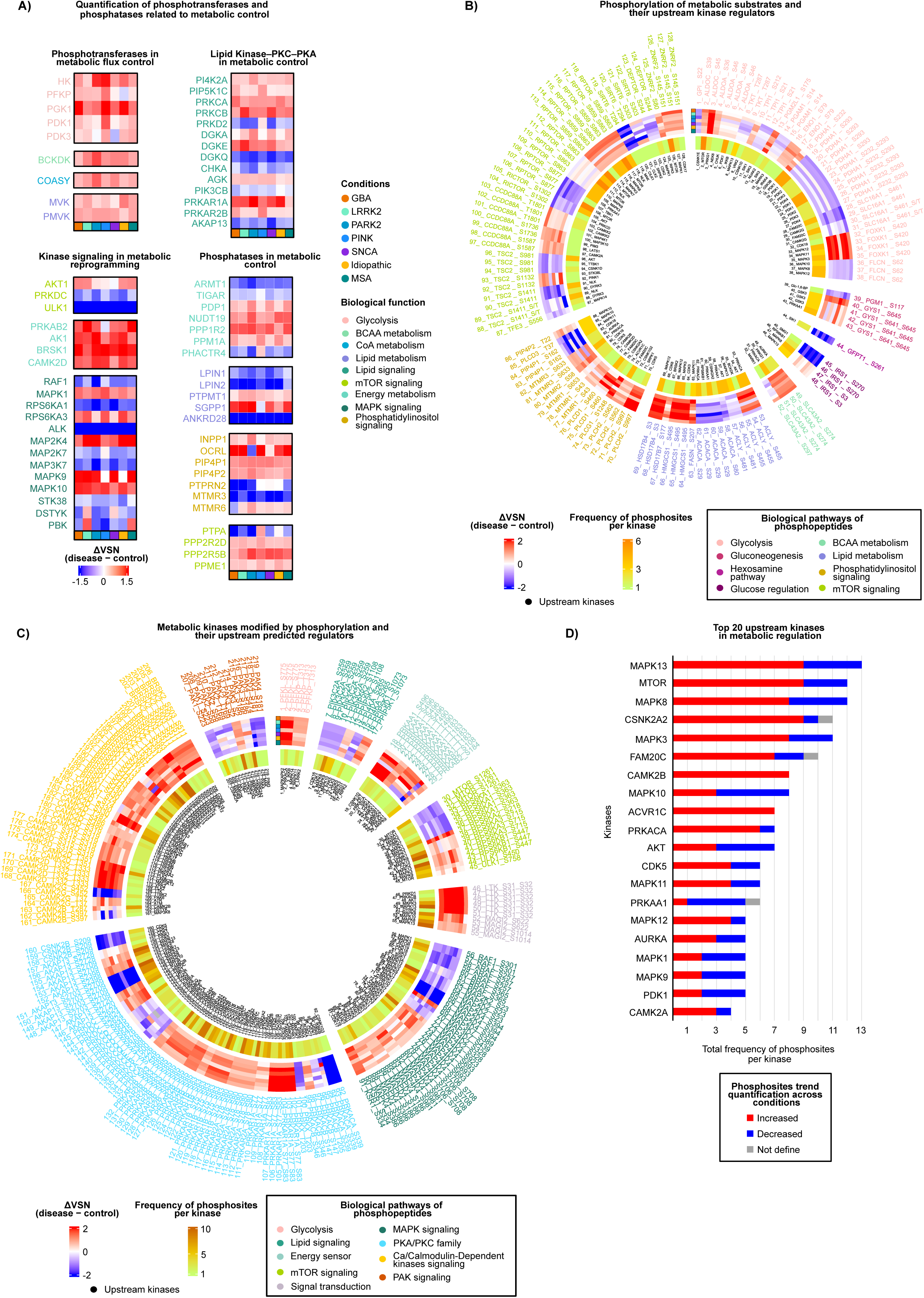
Phosphoproteomic analysis links metabolic dysfunction to altered kinase and phosphatase signaling in PD and MSA spheroids. (A) Quantification of phosphotransferases and phosphatases involved in metabolic control. Annotated heatmaps display differentially expressed and co-expressed phosphotransferases and phosphatases relative to controls (robust empirical Bayes, p < 0.05, |log2FC| > ±0.5; Mann–Whitney U test, p < 0.05). The major functional classes include regulators of metabolic flux and metabolic reprogramming, including components the lipid kinase–PKC–PKA signaling axis. (B–C) Regulatory phosphorylation landscape of metabolic proteins and their upstream kinase regulators. Annotated circular heatmaps display fold changes in phosphosite abundance for metabolic substrates (B) and phosphotransferases (C) involved in metabolic regulation relative to controls. The inner ring shows experimentally validated and computationally predicted upstream kinases regulating each phosphosite. (D) Top 20 upstream kinases associated with metabolic regulation in PD and MSA spheroids.

Kinase signaling modules associated with metabolic reprogramming and energy sensing were also remodeled at the protein abundance level. Disease spheroids showed increased levels of AKT1 and AMPK-associated subunits, such as PRKAB2, along with altered MAPK/ERK-linked nodes (e.g., MAPK1, RPS6KA1/3), consistent with broad shifts in nutrient and growth signaling. In contrast, reduced ULK1 and PRKDC levels suggest an imbalance between nutrient signaling and autophagy/stress response control (Figure 4A). Lipid second messenger signaling was prominently affected, with increased phosphoinositide and DAG-associated kinases and effectors (PI4K2A, PIP5K1C, DGKA/DGKE, AGK), together with increased PKC isoforms (PRKCA, PRKCB) and remodeling of PKA regulatory subunits, including PRKAR1A and PRKAR2B (Figure 4A). Conversely, reductions in regulators such as PRKD2, DGKQ, AKAP13, CHKA, and PIK3CB point to altered phosphatidylcholine synthesis and DAG turnover, linking phosphorylation signaling to membrane lipid remodeling. Phosphatase changes further reinforced this axis, including changes in PDP1, PTPMT1, NUDT19, LPIN1/2, and SGPP1, as well as broad remodeling of phosphoinositide phosphatases, including INPP1, OCRL, PIP4P1/PIP4P2, and MTMR family proteins. Together, these changes are consistent with coordinated rewiring of kinase and phosphatase control over metabolic and membrane pathways (Figure 4A).

We next examined phosphorylation-state remodeling directly by phosphoproteomics, mapping dysregulated phosphorylation events onto metabolic substrates and signaling pathways (Table S7-8). Pathway enrichment analysis of differentially phosphorylated peptides (Table S8) identified signal transduction as a dominant category across disease conditions, with consistent involvement of cAMP/PKA, AMPK, mTOR, phosphatidylinositol, and DAG/IP3-PKC/PLCβ modules, and MAPK family signaling, together with broad effects on membrane trafficking, ER-Golgi transport, autophagy, and apoptosis programs (Figure S2A). Phosphoproteomic WGCNA (Table S8) further revealed modules enriched for autophagy/AMPK signaling, immune-related programs, and membrane trafficking/ER-Golgi processes, supporting coordinated phosphorylation remodeling rather than isolated phosphosite changes (Figure S2B).

At the level of individual phosphosites, dysregulated phosphorylation encompassed glycolytic and glucose-regulatory enzymes, BCAA- and CoA-linked nodes, lipid biosynthesis and turnover enzymes, and phosphoinositide signaling proteins (Figure 4B). In parallel, multiple kinases also displayed altered phosphorylation states, revealing extensive feedback regulation across MAPK tiers, mTOR/energy-sensing nodes, Ca^2+^/calmodulin-dependent kinases, PAK family kinases, and second messenger-dependent kinases, including PKA and PKC (Figure 4C). Kinase-substrate prediction identified a restricted set of upstream drivers that dominated the metabolic phosphosite landscape, with strong representation of MAPK-family kinases and mTOR, together with contributions from CK2 (CSNK2A2), PKA (PRKACA), PDK1, AKT, CDK5, and CaMK-family kinases (Figure 4D). This architecture provides a mechanistic framework in which phosphorylation-dependent signaling networks may reinforce the glycolysis-mitochondria coupling defect, BCAA bottlenecks, and lipid/membrane remodeling observed in PD and MSA spheroids.

### 2.5. Conserved metabolic and phosphorylation-linked signaling signatures are detected in postmortem substantia nigra from PD and MSA donors

To determine whether spheroid-defined molecular axes reflect disease-relevant processes in vivo, we analyzed postmortem brain tissue (PMBT) from substantia nigra samples obtained from controls (n = 4), individuals with idiopathic PD (n = 4), and individuals with MSA (n = 6) using the same multi-omics framework ((Table S1; Table S9-10; Figure S1). Global proteomic PCA of PMBT separated control, PD, and MSA samples more distinctly than in spheroids, consistent with stronger disease-specific divergence in end-stage human tissue (Figure S3A). Marker profiling showed changes in dopaminergic-associated proteins (TH, DDC), together with oligodendroglial/myelin-associated proteins (MBP, PLP, CNP), supporting both nigral dopaminergic and myelin-related pathology while also indicating disease-associated differences in cellular composition (Figure S3B). Protein groups linked to dopamine synapse biology, mitochondrial function, ubiquitin-proteasome activity, membrane trafficking, and autophagy/lysosome pathways were also altered (Figure S3C). Dopamine-related metabolites showed disease-associated alterations, including changes in tyrosine/phenylalanine-linked metabolites and dopamine catabolites (Figure S3D). Dopamine itself was reduced in PD and undetectable in MSA, which may reflect very low abundance caused by severe nigral cell loss, greater in MSA than PD, rather than absence of dopamine-related biology.

Even with greater global separation between PD and MSA in PMBT, metabolic processes again emerged as a dominant shared alteration. Across differentially regulated proteins in PMBT, metabolic pathways accounted for ∼51% of dysregulated proteins (Figure 5A). Enrichment of metabolism-related proteins emphasized lipid-centered programs, particularly glycerophospholipid biosynthesis and broader phospholipid metabolism, together with additional metabolic categories (Figure 5A).

**Figure 5.**
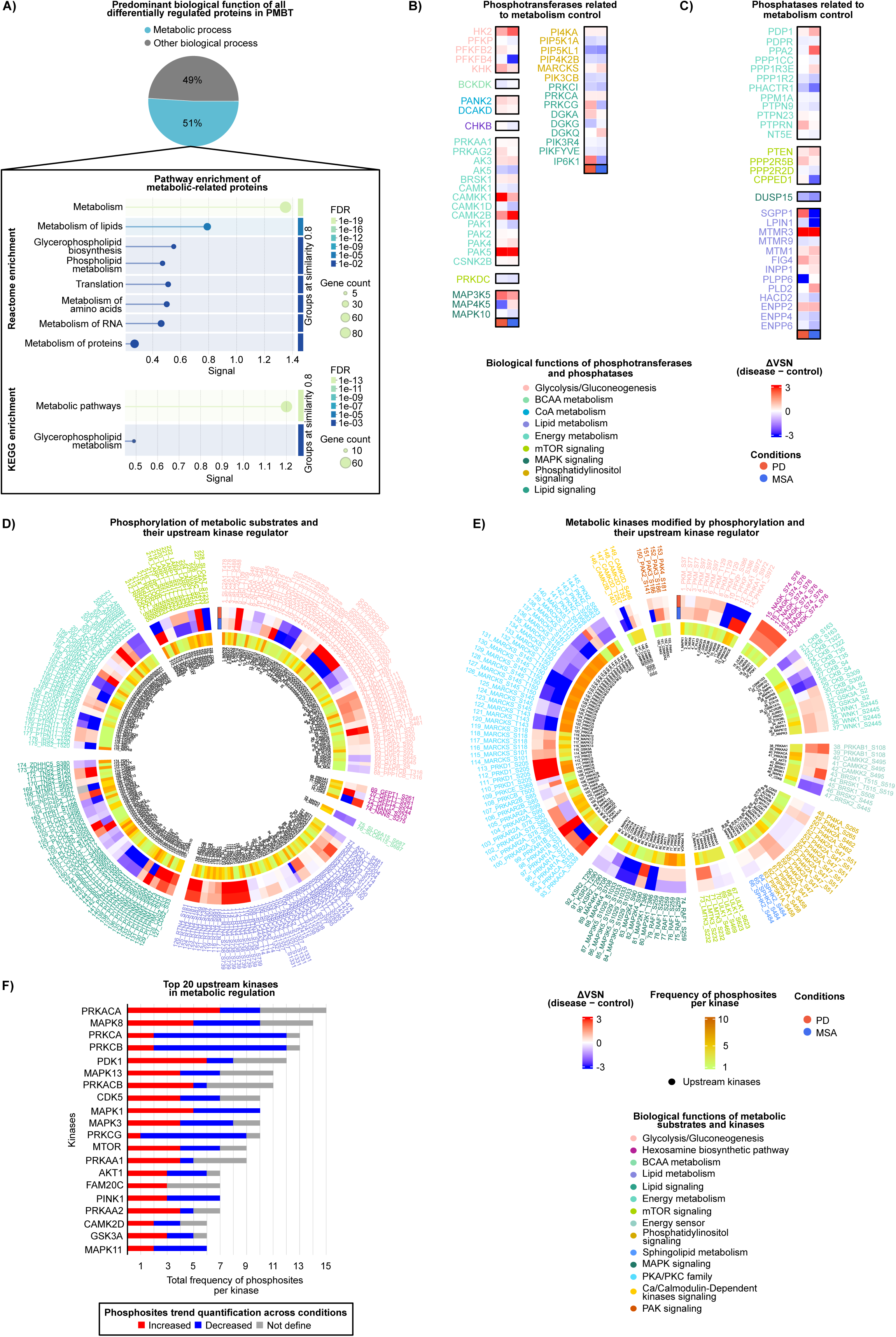
Postmortem substantia nigra proteomics and phosphoproteomics reveal conserved metabolic signaling alterations in PD and MSA. (A) Approximately 51% of proteins dysregulated in substantia nigra postmortem brain tissue (PMBT) from patients with PD and MSA are associated with metabolic processes. Pathway enrichment analysis was performed to refine the metabolic category into specific functional pathways. (B–C) Abundance profiling of phosphotransferases (B) and phosphatases (C) involved in metabolic regulation. Annotated heatmaps display differentially expressed and co-expressed phosphotransferases and phosphatases in PD and MSA PMBT relative to control PMBT (robust empirical Bayes, p < 0.05, |log2FC| > ±0.5; Mann–Whitney U test, p < 0.05). (D–E) Phosphorylation landscape of metabolic proteins and their upstream kinase regulators in substantia nigra PMBT. Circular heatmaps summarize phosphosite fold changes across metabolic substrates (D) and phosphotransferases (E), with experimentally validated and computationally predicted upstream kinases regulating each phosphosite shown in the inner ring. (F) Top 20 upstream kinases associated with metabolic regulation in PD and MSA PMBT.

To define shared architecture across molecular layers in PMBT, we integrated pathway and network analyses across proteomics, metabolomics, and phosphoproteomics (Table S10). Integrated multi-omics analysis revealed convergence between PD and MSA in metabolic pathways, including glucose/carbon metabolism, amino acid metabolism (including BCAAs), CoA-related processes, and lipid remodeling, alongside proteostasis, trafficking, and immune signaling pathways (Figure S4A-D). Proteomic WGCNA identified modules distinguishing controls from PD/MSA samples (Table S10), with enrichment for protein metabolism, oxidative phosphorylation, metabolic and immune signaling pathways, including IFN-stimulated genes and interleukin-family signaling, autophagy, Rho GTPase programs, and sphingolipid signaling (Figure S4B). These findings reinforce the prominence of metabolic remodeling embedded within broader signaling and trafficking architecture.

At the level of shared metabolic components, PMBT displayed coordinated changes in proteins linked to glycolysis and glycogen metabolism (e.g., GPI, GAPDH, PYGL, AGL), mitochondrial pyruvate import and TCA-linked enzymes (e.g., MPC1/2), PDHB, OGDHL, ME1), and the hexosamine/serine biosynthesis interface (e.g., OGT, PHGDH) (Figure S4E). BCAA-associated enzymes (e.g., BCAT2, BCKDHB, IVD, PCCA, MMUT) and CoA-linked nodes (including pantothenate transport and acyl-CoA handling such as SLC5A6 and ACOT1) were also dysregulated, supporting a conserved BCAA-CoA bottleneck axis in human tissue. Lipid metabolic alterations encompassed fatty-acid transport/oxidation (e.g., ABCD1/2, CPT1A, ACADVL/ACADM), lipogenesis and citrate/acetyl-CoA routing (e.g., SLC25A1, FASN), sterol metabolism (e.g., HMGCS1, DHCR7), and membrane lipid remodeling and signaling proteins, including PLA2G4C, SMPD1, and phosphoinositide-linked proteins (Figure S4E).

Metabolomics reinforced these protein-level observations in PMBT, including signatures consistent with restricted glycolytic entry and redox rerouting, such as reduced glucose and glucose-6-phosphate with increased sorbitol, glycerol-3-phosphate, and α-hydroxybutyrate. Altered mitochondrial flux, changes in citrate and downstream TCA intermediates, and BCAA-related disruption also were observed, including altered BCAA levels with reduced downstream catabolic products and propionylcarnitine-linked features. These changes occurred together with broad lipid remodeling, including altered fatty acids, lysophospholipids, and phosphoinositide-related metabolites (Figure S4F).

Finally, phosphorylation-centered analyses indicated that these conserved metabolic states are coupled to rewired kinase-phosphatase control in PMBT. Profiling of phosphotransferases and phosphatases associated with metabolic regulation revealed remodeling of glycolytic entry/commitment nodes (e.g., HK2, PFKP, PFKFB2/4, KHK), BCAA/CoA-linked regulators (e.g., BCKDK, PANK2, DCAKD), and lipid second messenger signaling components, including phosphoinositide and DAG regulators such as PI4K4, PIP5K1A, and PGK family proteins (Figure 5B and C). Phosphorylation signatures spanning metabolic substrates and kinases recapitulated a membrane- and stress-kinase-centered signaling architecture (Figure 5D and E), and upstream kinase inference again highlighted dominant contributions from PKA/PKC-family kinases and MAPK family kinases, with additional nutrient/energy-sensing regulators including mTOR, AKT, PDK1, and AMPK-related nodes (Figure 5F). Together, these findings support the conservation of a phosphorylation-linked metabolic axis between patient-derived spheroids and human substantia nigra, while capturing the increased disease-specific divergence observed in postmortem tissue.

### 2.6. The metabolic landscape perturbed in PD and MSA: from metabolic substrates to kinase-network regulation

To evaluate the extent to which the metabolic and signaling programs identified in iPSC-derived midbrain spheroids are conserved in human disease tissue, we performed a cross-system concordance analysis between spheroids and PMBT (Table S11). For each dataset, disease-associated changes in PD and MSA versus controls were compared across sample types using median ΔVSN disease-control differences calculated separately for each condition and sample type. Features were classified as shared or divergent based on their quantitative behavior (Figure S6). This analysis showed that the core PD/MSA metabolic signature observed in spheroids shows substantial concordance with PMBT. Together, this framework enabled us to build an integrated map of the perturbed metabolic axes across these alpha-synucleinopathies, linking altered metabolites and proteins to phosphosite-level regulation and predicted upstream kinase networks (Figure 6).

**Figure 6.**
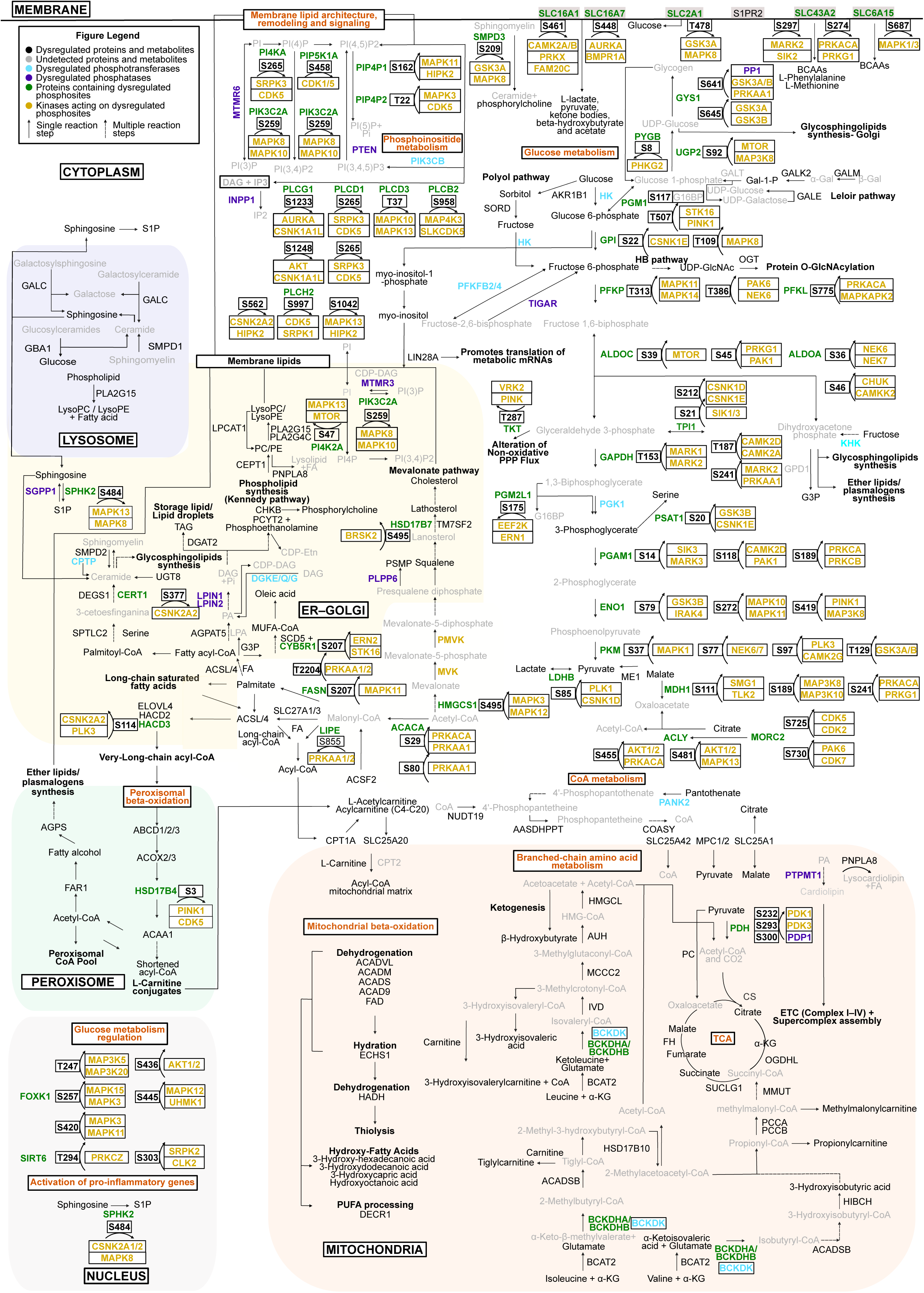
Integrated multi-omics mapping defines convergent metabolic dysfunction across spheroid models and postmortem brain tissue. Integrated mapping of convergent metabolic dysfunction across spheroids and PMBT in PD and MSA. All dysregulated metabolites, proteins, and phosphosites, together with the top predicted upstream kinases targeting metabolism-associated phosphosites in at least one condition, were incorporated into the integrative analysis.

Glucose metabolism, the TCA cycle, membrane lipid metabolism, mitochondrial and peroxisomal β-oxidation, BCAA metabolism, phosphoinositide signaling, and membrane lipid architecture/remodeling formed the central scaffold of metabolic perturbation across PD and MSA. Rather than representing isolated alterations, the integrated map revealed these axes as an interconnected network spanning the cytoplasm, mitochondria, peroxisomes, ER-Golgi system, lysosomes, plasma membrane, and nucleus. Within this scaffold, glucose metabolism emerged as a central phosphorylation-regulated axis, with phosphosites and predicted upstream kinases distributed across nearly all steps of glycolysis and pyruvate handling. This suggests that glycolysis and pyruvate metabolism occupy a central regulatory position, coordinating carbon allocation between ATP production, mitochondrial substrate entry, redox balance, biosynthetic routes, and membrane lipid remodeling.

This organization also provides a potential mechanistic link among the conserved alterations in BCAA metabolism, pantothenate/CoA-linked metabolites, lysophospholipids, and sphingolipid-related species observed in both spheroids and PMBT. BCAA catabolism, fatty acid activation, β-oxidation, acetyl-CoA production, acylcarnitine formation, and lysophospholipid reacylation all depend on CoA or acyl-CoA availability. Therefore, changes in pantothenate, a precursor of CoA biosynthesis, suggest pressure on the CoA/acyl-CoA economy, potentially creating a bottleneck between mitochondrial carbon utilization to lipid remodeling. Altered lysophospholipids may reflect disrupted membrane repair and reacylation cycles, whereas sphingolipid and glycosphingolipid changes may arise from altered palmitoyl-CoA-dependent ceramide synthesis, lysosomal lipid handling, and galactose-Leloir/sugar-nucleotide metabolism. Together, these findings support a model in which PD and MSA share a conserved metabolic-regulatory program that couples glucose/pyruvate control, CoA-dependent substrate utilization, BCAA catabolism, and membrane lipid remodeling across cellular compartments.

This integrative landscape (Figure 6) is organized by an upstream kinase network (Figure S7), acting in concert with metabolic signals such as cAMP, AMP, ADP, and leucine. We delineated the major signaling modules positioned to regulate the metabolic state. These modules include phosphoinositide/DAG-PKC signaling at the membrane, insulin/IGF-AKT signaling, ERK/MAPK and JNK stress-responsive pathways, mTORC1-autophagy-lysosome regulation, mTORC2/ER-associated signaling, AMPK energy sensing, CaMK/Ca²⁺-dependent signaling, and nucleotide/cAMP-PKA signaling. Together, this map places the phosphorylation changes observed in PD and MSA within a broader regulatory framework that senses metabolic imbalance and translates it into coordinated kinase-driven cellular responses. Rather than forming independent pathways, these kinase modules are highly interconnected, with multiple shared regulatory nodes converging not only on metabolic control and lipid signaling but also on other cell functions in a coordinated manner, including autophagy, nutrient sensing, lysosomal function, calcium responses, and proteostasis.

### 2.7. Metabolic-kinase rewiring converges on aSYN post-translational regulation

The conserved metabolic and phosphorylation-linked signaling alterations observed across PD and MSA were associated with cellular processes directly relevant to synucleinopathy, including autophagy, lysosomal function, membrane trafficking, and proteostasis. We therefore asked whether the kinase networks remodeled in midbrain spheroids and postmortem substantia nigra could also intersect with post-translational modification sites on aSYN that have been implicated in oligomerization and aggregation. To address this, we performed kinase-substrate mapping for major aSYN phosphosites and compared the predicted aSYN-targeting kinases with dysregulated and phosphosite-predicted kinases detected in spheroids and PMBT. This analysis identified kinase groups selectively represented in spheroids or PMBT, as well as a shared kinase set across both systems (Figure 7A).

**Figure 7.**
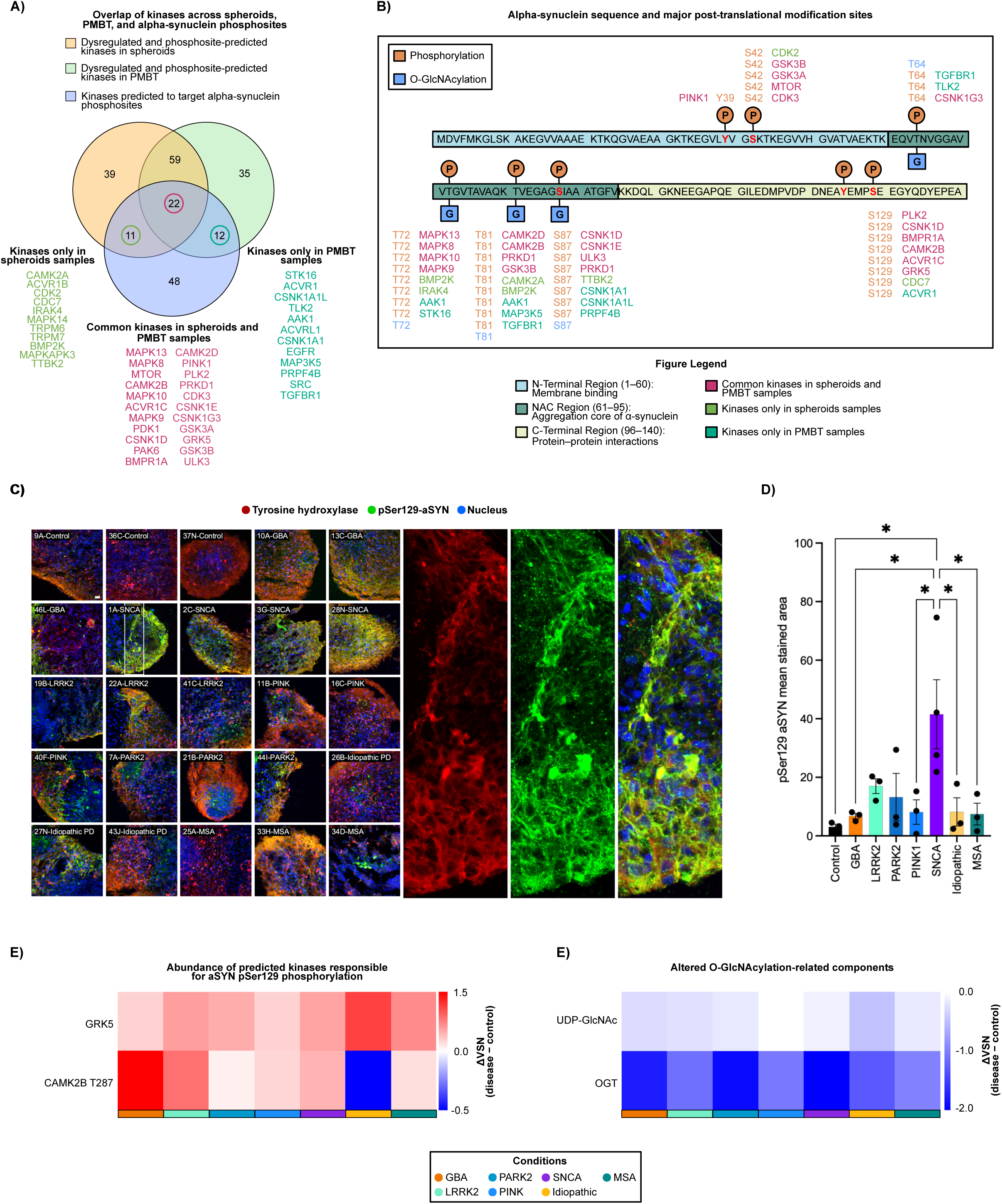
Metabolism-linked kinase signaling connects α-synuclein phosphorylation, O-GlcNAcylation, and pathological aggregation in PD and MSA. (A) Overlap of kinases across spheroids, PMBT, and alpha-synuclein phosphosites. Dysregulated and phosphosite-inferred kinases associated with metabolic regulation in spheroids and PMBT are also predicted to target α-synuclein phosphosites. Venn diagram highlights both dataset-specific kinase subsets and a substantial shared kinase pool. (B) α-synuclein sequence and major post-translational modification sites. Schematic representation of the SNCA amino acid sequence highlighting site-specific predicted kinases identified in (A). These kinases target key residues that are also subject to O-GlcNAcylation, a nutrient- and metabolism-sensitive post-translational modification that functionally cross-talks with phosphorylation. O-GlcNAc modification of SNCA has been implicated in regulating its aggregation. The overlap between kinase-targeted phosphosites and O-GlcNAcylation sites suggests a potential mechanistic link between metabolic perturbations and pathological SNCA aggregation in PD and MSA. (C) Representative immunofluorescence images of midbrain spheroids stained for DAPI (blue), pSer129-aSYN (green) and tyrosine hydroxylase (red). Panels on the right show high magnification views of the boxed inset from line 1A-SNCA. (D) Quantification of pSer129-aSYN signal in midbrain spheroids, showing increased pSer129-aSYN in disease spheroids. Statistical significance was determined by one-way ANOVA, * p < 0.05. scale bar is 20 mm. (E) Abundance of predicted kinases responsible for aSYN pSer129 phosphorylation. The predicted kinases GRK5 and CAMK2B T287 were detected as dysregulated in or datasets (robust empirical Bayes, p < 0.05, |log2FC| > ±0.5) (F) Altered O-GlcNAcylation-related components. UDP-GlcNAc and OGT shows a significant reduction in our datasets (OPLS classification cutoff > 0.75; (robust empirical Bayes, p < 0.05, |log2FC| > ±0.5).

Mapping these kinases onto the aSYN sequence showed predicted regulation of multiple aggregation-relevant residues, including sites within or near the NAC aggregation-prone region, such as T64, T72, and T81, as well as C-terminal residues (S87 and S129) (Figure 7B). Notably, S129 was predicted to be targeted by several disease-associated kinases, including GRK5, CAMK2B, PLK2, CSNK1D, BMPR1A, ACVR1C, and ACVR1. Consistent with this prediction, disease spheroids showed an increased trend in pS129-aSYN phosphorylation (Figure 7C and D), concomitant with increased levels of GRK5 and phosphorylated CAMK2B T287 in almost all conditions (Figure 7E).

Several SNCA residues predicted to undergo phosphorylation also overlap with, or lie close to, reported O-GlcNAcylation sites. O-GlcNAcylation at residues such as T72^36^ and T81^37^ has been associated with reduced aSYN aggregation. Interesting, our data also shows a significant reduction of in O-GlcNAc-related components such as UDP-GlcNAc and OGT (Figure 7F), suggesting a potential altered balance between phosphorylation and O-GlcNAcylation at aggregation-sensitive residues. Together, these findings suggest a mechanistic link among metabolic dysfunction, kinase-network remodeling, and aSYN post-translational regulation.

## 3. Discussion

Parkinson’s disease (PD) and multiple system atrophy (MSA) are alpha-synucleinopathies with overlapping clinical manifestations but distinct cellular patterns of pathology. Although pathological aSYN accumulation defines both disorders, the molecular programs that accompany and potentially shape PD and MSA extend beyond protein aggregation. Our findings support the concept that metabolic dysfunction represents a central disease-associated program connecting these alpha-synucleinopathies. Metabolic perturbation emerged as the dominant molecular alteration shared by PD and MSA, accounting for the largest proportion of disease-associated changes and showing convergence across patient-derived iPSC midbrain spheroids and postmortem substantia nigra. Altered cellular metabolism has been consistently observed in PD, MSA, and other neurodegenerative disorders, where it is increasingly recognized as a contributor to cellular vulnerability, rather than a passive consequence of degeneration^38–40^. Importantly, despite differences in cellular composition, maturation state, and pathological complexity between patient-derived spheroids and PMBT, detecting these metabolic-regulatory programs in both systems supports their relevance to PD and MSA pathology.

This convergent metabolic axis was organized around an interconnected glucose–TCA–BCAA–CoA–lipid remodeling program. In both PD and MSA, glucose and pyruvate metabolism occupied a central regulatory position, linking carbon entry to mitochondrial oxidation, routing of carbon into biosynthetic pathways, redox buffering, nucleotide sugar metabolism, and lipid synthesis. Impaired coupling between glycolysis and the TCA cycle may increase reliance on alternative energetic substrates, including BCAAs and lipids, to support TCA-cycle input and energy production. However, efficient use of these substrates depends on the availability of free CoA and the generation of multiple acyl-CoA intermediates. Accordingly, our data also show concomitant alterations in metabolites and proteins related to pantothenate/CoA metabolism, suggesting increased pressure on CoA availability and acyl-CoA handling. This provides a mechanistic link between impaired mitochondrial substrate utilization and lipid remodeling, including changes in lysophospholipids, sphingolipids, glycosphingolipids, and galactose-Leloir/sugar-nucleotide pathways. These results are consistent with previous studies reporting glucose metabolic impairment^41^ and hypometabolism^42^, mitochondrial dysfunction^20^, BCAA accumulation^43,44^ and profound lipid remodeling^45,46^ in PD and other neurodegenerative disorders. Together, these findings suggest that PD and MSA share an integrated metabolic state in which altered carbon allocation, CoA-dependent substrate utilization, and membrane lipid remodeling converge as coordinated manifestations of a common metabolic constraint.

Kinase-substrate analyses indicate that this metabolic disturbance is embedded within a broader phosphorylation-responsive regulatory architecture. Phosphotransferases and phosphatases linked to glycolytic commitment, pyruvate-to-acetyl-CoA conversion, BCAA flux, CoA metabolism, and phosphoinositide/DAG-centered lipid signaling were altered across PD and MSA, positioning phosphorylation as a regulatory layer over the shared metabolic axis. In parallel, AMPK, mTOR, MAPK/JNK, PKA/PKC, CaMK, and insulin/AKT signaling modules connect energetic state, nutrient availability, membrane lipid composition, calcium signaling, and stress responses. Several of these pathways have previously been associated with PD^47–51^ and neurodegenerative disorders. Our results therefore suggest that metabolic dysfunction in PD and MSA may represent more than a downstream consequence of disease, but part of a coordinated kinase-regulated state capable of sensing, reinforcing, or adapting to metabolic bottlenecks. In particular, convergent changes in AMPK/mTOR nutrient sensing, MAPK/JNK stress signaling, and lipid second-messenger pathways provide a regulatory bridge between metabolism and other dysfunctional processes previously reported in PD and MSA, including autophagy, lysosomal function, inflammation, membrane trafficking, and proteostasis.

The metabolic-kinase axis identified here also provides a potential link between metabolic dysfunction and aSYN post-translational regulation. Several kinases predicted to target aSYN phosphosites, including S129 and residues within or near aggregation-relevant regions, were dysregulated or inferred from phosphoproteomic analyses in spheroids and postmortem substantia nigra. In parallel, our integrated metabolic map identified alterations in O-GlcNAc-related components, including UDP-GlcNAc and OGT, within the broader glucose and hexosamine-pathway disturbance observed across PD and MSA. Because some predicted phosphorylation sites overlap with, or lie close to, reported O-GlcNAcylation sites, these findings suggest that impaired glucose metabolism may influence aSYN not only through bioenergetic stress, but also by altering the balance between phosphorylation and O-GlcNAcylation at aggregation-sensitive residues. This is relevant because O-GlcNAcylation at specific aSYN residues has been associated with reduced aggregation, whereas increased pS129-aSYN is a prominent feature of synucleinopathy pathology. Although kinase-substrate relationships require experimental validation, the overlap among dysregulated metabolic kinases, altered O-GlcNAc-related metabolism, and increased pS129-aSYN suggests that PD and MSA may converge on a shared regulatory axis linking metabolic dysfunction, kinase-network remodeling, and aSYN modification states that promote aggregation-prone conformations.

The metabolic-regulatory disturbance identified here may represent an underlying cellular state that facilitates or amplifies vulnerability, even when disease arises from distinct genetic or sporadic contexts. This is particularly relevant because the PD spheroids included in this study capture distinct molecular routes into disease, including PINK1- and PRKN/PARK2-associated mitochondrial quality-control defects^52^, GBA-linked lysosomal dysfunction^53^, and LRRK2-associated vesicular and kinase signaling alterations^54^, SNCA dosage-related pathology caused by SNCA duplications^55^ or triplications^56^, and idiopathic forms. These genetic backgrounds differ in penetrance, suggesting that additional cellular states may influence disease vulnerability. This idea is also consistent with epidemiological studies linking idiopathic PD to systemic metabolic dysfunction, including type 2 diabetes, insulin resistance, and metabolic syndrome^57–59^. In this context, the metabolic-regulatory axis identified here may influence disease risk or progression by affecting substrate handling, lipid remodeling, proteostasis, and aSYN post-translational regulation. Collectively, this shared vulnerability state highlights metabolic regulation as a potential point of intervention across synucleinopathies.

Several components of the metabolic perturbation identified in PD and MSA resemble metabolic signatures associated with type 2 diabetes, insulin resistance, and metabolic syndrome. These include altered glucose handling^60,61^, BCAA accumulation^62,63^, lipid dysregulation^64,65^, and changes in metabolic-kinase networks. This does not imply that PD or MSA are systemic metabolic diseases but rather suggests that synucleinopathies may engage conserved cellular programs that are also prominent in metabolic disorders. Consistent with this idea, several metabolism-targeting strategies, including GLP-1 receptor agonists^66^, insulin-sensitizing PPARγ agonists^67^, mitochondrial bioenergetic modulators such as CoQ10/ubiquinol^68,69^ and UDCA^70^, and urate-elevating strategies^71^, have been tested in synucleinopathies with mixed clinical outcomes. Existing interventions have largely targeted glucose/insulin signaling, mitochondrial bioenergetics, redox metabolism, or lysosomal lipid handling. Our data extend this therapeutic framework by identifying a CoA-linked metabolic axis that integrates glucose-pyruvate utilization, BCAA catabolism, and lipid remodeling. Future strategies aimed at restoring metabolic balance in synucleinopathies may therefore benefit from targeting broader metabolic regulatory nodes rather than single downstream pathways.

## 4. Limitations

Although our integrative analyses reveal a conserved metabolic architecture across PD and MSA, several limitations should be considered. First, iPSC-derived midbrain spheroids provide a powerful platform for modeling human dopaminergic neurons, but they do not fully recapitulate the cellular diversity, vascularization, aging, or multicellular architecture of the human brain. In particular, these models lack mature oligodendrocytes and the complex neuron-glia interactions, which are especially relevant for MSA pathology. Second, analyses were performed at day 100 of differentiation, representing a relatively early maturation stage that cannot capture the long-term, age-associated processes that unfold over decades in PD and MSA. At the same time, because iPSC-derived brain spheroids (like organoids) are a model for studying early brain development, and because reprogramming rests aging-associated features while generating cells with embryonic-like properties^72–77^, these models raise an important conceptual question: whether disease-relevant cellular dysfunction may originate during central nervous system development, when genetic programs and regulatory networks are first established, but remain compensated and clinically silent for decades until aging and systemic metabolic changes amplify these latent vulnerabilities, driving pathology and the eventual emergence of clinical symptoms. Third, although the study included familial and idiopathic PD lines as well as MSA patient-derived models, the number of lines per subgroup was modest, and larger cohorts will be required to capture the full spectrum of genetic and clinical heterogeneity. Finally, while this approach identifies convergent metabolic programs across experimental models and human tissue, it does not fully resolve the temporal ordering of these alterations relative to disease initiation and progression. Despite these limitations, convergence of protein, metabolite, and phosphorylation-linked signaling changes across iPSC-derived midbrain spheroids and post-mortem human brain tissue identifies metabolic-kinase remodeling as a central and shared molecular hallmark of PD and MSA.

## 5. Experimental procedures

### 5.1. Patient identification, protection, sampling and iPSC lines generation

All procedures were conducted I accordance with Swedish national and European Union directives. The patient biopsies utilized to generate the iPSCs were obtained with informed consent and after ethical committee approval at the Parkinson Institute in Milan, Italy (Ethics Committee “Milano Area C” and registered under the number 370-062015. The permit for reprogramming was delivered by the Swedish work environment authority to L.R. registered under the number 20200-3211. The generation of the iPSC lines was previously published^78–90^.

### 5.2. Generation of iPSC-derived midbrain spheroids

iPSC-derived midbrain spheroids were generated using a ventral midbrain dopaminergic differentiation protocol adapted from Chumarina et al.^91^ Briefly, following expansion, iPSC colonies were detached with Dispase II and transferred to ultra-low-attachment culture flasks in WiCell medium supplemented with Y-27632 ROCK inhibitor and 20 ng/mL FGF2. The following day was defined as day 0 of differentiation. Medium was replaced with neural induction medium consisting of Advanced DMEM/F12 supplemented with 2 mM L-glutamine, 1% non-essential amino acids, 1% N2 supplement, and 1% penicillin–streptomycin. To induce neuralization and ventral midbrain patterning, cultures were treated from days 0–4 with 0.1 µM LDN-193189, 10 µM SB-431542, 200 ng/mL SHH-C, 1 µM SAG, and 0.8 µM CHIR-99021, with medium changes every other day. On day 6, SB-431542 and SHH-C were withdrawn, and SAG was increased to 2 µM. LDN-193189 was removed on day 10. From day 12 onward, spheroids were maintained in neural induction medium supplemented with 100 ng/mL FGF8, 2 µM SAG, 10 ng/mL BDNF, and 200 µM ascorbic acid. On day 22, medium was changed to neural differentiation medium composed of Neurobasal medium supplemented with 2 mM L-glutamine, 1% non-essential amino acids, 1% N2 supplement, 1% B27 without vitamin A, and 1% penicillin–streptomycin. The medium was further supplemented with 100 ng/mL FGF8, 2 µM SAG, 10 ng/mL BDNF, 10 ng/mL GDNF, 200 µM ascorbic acid, 500 µM dibutyryl-cAMP, and 1 ng/mL TGF-β. From day 30 onward, FGF8 and SAG were removed. 50 µM of dopamine was added during later maturation to promote neuromelanin-associated maturation. Spheroids were maintained until day 100 in vitro, when they were washed and collected for downstream proteomic, phosphoproteomic, metabolomic, and immunostaining analyses.

### 5.3. iPSC-derived midbrain spheroid staining

For immunocytochemistry, day-100 iPSC-derived midbrain spheroids were fixed in 4% paraformaldehyde, washed in PBS, equilibrated overnight in 30% sucrose, embedded in OCT compound, and cryo-sectioned at 20 µm thickness. Spheroid sections were blocked in PBS containing 10% donkey serum. Dissociated cells were permeabilized with 0.1% Tween-20, whereas cryosections were permeabilized with 0.3% Triton X-100. Primary antibodies were diluted in blocking solution and incubated overnight at 4 °C. The staining panel included antibodies against TH, FOXA2, MAP2, and pS129-aSYN. After washing, samples were incubated for 1 h at room temperature with Alexa Fluor-conjugated secondary antibodies diluted 1:400 in PBS. Nuclei were counterstained with DAPI. Images were acquired using an inverted epifluorescence microscope, and image analysis and quantification were performed using MetaMorph (Molecular Devises).

### 5.4. Collection of PD and MSA post-mortem brain tissue

Postmortem substantia nigra samples were obtained post-mortem from individuals with Parkinson’s disease (PD, n = 4), multiple system atrophy (MSA, n = 6), and neurologically healthy controls (n = 4) through the Netherlands Brain Bank. Ethical approval for the use of these tissues was granted by the regional review board in Malmö-Lund, Sweden (under permit numbers M46-16 and M69-16).

### 5.5. Sample preparation

#### 5.5.1. Metabolomic analysis

Metabolic profiling of spheroids and postmortem substantia nigra tissue was carried out at the Swedish Metabolomics Centre (SciLifeLab, Umeå, Sweden). Approximately 20 mg of brain tissue was extracted using our well-established protocol^92^. Briefly, samples were homogenized in 1,000 µL of extraction buffer (80:20 methanol:water, v/v) containing internal standards for GC-MS and LC-MS. Homogenization was performed with a tungsten bead at 30 Hz for 3 min, after which the bead was removed and the samples centrifuged at 14,000 rpm (18,620 g) for 10 min at 4 °C. Aliquots of the supernatant (200 µL for LC-MS, 50 µL for GC-MS) were dried in a vacuum concentrator and stored at −80 °C until analysis. LC-MS/MS runs were additionally performed on two pooled samples for compound identification. Samples were randomized before analysis on both platforms.

#### 5.5.2. Proteomic and phosphoproteomic analyses

Proteins were isolated from day 100 iPSC-derived midbrain spheroids using a lysis buffer containing 25 mM DTT and 10% (w/v) SDS in 100 mM triethylammonium bicarbonate (TEAB). Lysates were disrupted by sonication (40 cycles, 15 s on/15 s off, 4 °C; Bioruptor Plus, UCD-300, Diagenode), followed by heating at 99 °C for 5 min. Samples were then centrifuged at 20,000 × g for 15 min at 18 °C, and protein concentrations in the collected supernatants were determined with the Pierce 660 nm Protein Assay in combination with the Ionic Detergent Compatibility Reagent. Extracted proteins were stored at −80 °C until further processing.

Protein digestion was performed using the S-Trap™ 96-well plate method (ProtiFi) according to the manufacturer’s protocol. Briefly, proteins were alkylated with 50 mM iodoacetamide (IAA) for 30 min at room temperature in the dark, followed by overnight trypsin digestion at 37 °C in 50 mM TEAB using an enzyme-to-substrate ratio of 1:50. Peptides were sequentially eluted with 80 µL 50 mM TEAB, 80 µL 0.2% formic acid (FA), and 80 µL 50% acetonitrile (ACN) containing 0.2% FA. Eluates were dried using a vacuum concentrator and resuspended in 0.1% trifluoroacetic acid (TFA) and 2% ACN. Peptide concentrations were quantified using the Pierce Quantitative Colorimetric Peptide Assay. One microgram of peptides was injected into the LC-MS/MS system for further global proteomic analysis. For phosphoproteomic profiling, 55 µg of peptides underwent C18-based cleanup on the automated AssayMAP Bravo system (Agilent Technologies). Following desalting, phosphopeptides were isolated by Fe(III)-IMAC affinity chromatography as described previously. Enriched fractions were then resuspended in 2% acetonitrile (ACN) with 0.1% trifluoroacetic acid (TFA) and analyzed by nanoLC-MS/MS.

### 5.6. Mass spectrometry analysis

#### 5.6.1. GC-MS analysis of metabolites

Derivatization and GC-MS acquisition were performed as previously described^93^. Dried extracts were derivatized with 30 µL methoxyamine (15 µg/µL in pyridine) followed by 60 µL of MSTFA (+ 1% TMCS) in heptane containing methyl stearate (15 ng/µL). One microliter of the derivatized solution was injected in splitless mode using an L-PAL3 autosampler (CTC Analytics, Switzerland) into an Agilent 7890B GC fitted with a 10 m × 0.18 mm, 0.18 µm Rxi-5 Sil MS column (Restek, USA). The injector was set to 270 °C, with purge activated after 60 seconds at a flow of 20 mL/min. The oven temperature was held at 70 °C for 2 min, increased at 40 °C/min to 320 °C, and maintained for 2 min. Detection was performed with a Pegasus BT GC-TOFMS (LECO, USA), using a 70 eV ionization beam, an ion source temperature of at 200 °C, a transfer line temperature of 250 °C, and an acquisition range of m/z 50–800 at 30 spectra/s.

#### 5.6.2. LC-MS analysis of metabolites

For LC-MS analysis, dried extracts were reconstituted in 10 µL methanol and 10 µL water. Each sample was analyzed in both positive and negative ionization modes on an Agilent 1290 Infinity UHPLC coupled to an Agilent 6546 Q-TOF mass spectrometer. Chromatographic separation was achieved on an Acquity UPLC HSS T3 column (2.1 × 50 mm, 1.8 µm) with a VanGuard pre-column, maintained at 40 °C. Mobile phases were (A) water with 0.1% formic acid and (B) 75:25 acetonitrile:2-propanol with 0.1% formic acid, at a flow of 0.5 mL/min. The gradient increased from 0.1% to 10% B over 2 minutes, ramped to 99% B over 5 minutes, held for 2 minutes, and re-equilibrated before the next injection. Electrospray ionization was performed in jet stream mode with identical settings across polarities, except for the capillary voltage. Continuous mass calibration was achieved by infusion of purine (m/z 121.05, 119.0363) and HP-0921 (m/z 922.0098, 966.0007). Data were collected over m/z 70–1700 at 4 scans/s.

#### 5.6.3. nanoLC-MS/MS analysis for proteomics

For nanoLC–MS/MS analysis, peptides were analyzed on an Exploris 480 mass spectrometer coupled to a Vanquish Neo nano UPLC system (Thermo Scientific) with an EASY-Spray ion source. Samples were run in randomized order using high-resolution data-independent acquisition (HR-DIA). Peptides were loaded onto an Acclaim PepMap 100 C18 trap column (75 µm × 2 cm, 3 µm, 100 Å, nanoViper) and separated on an Acclaim PepMap RSLC C18 analytical column (75 µm × 50 cm, 2 µm, 100 Å) at a flow rate of 300 nL/min and column temperature of 60 °C. Separation was achieved using a 120-min nonlinear gradient with mobile phases consisting of (A) 0.1% formic acid and (B) 0.1% formic acid in 80% acetonitrile. Solvent B was increased from 5% to 25% over 100 min, then to 32% over 12 min, to 45% over 8 min, and finally to 95% in 2 min, which was held for an additional 13 min. Mass spectrometry data were acquired in HR-DIA mode with a complete acquisition cycle consisting of 3 full MS1 scans followed by 18 MS2 DIA scans using variable isolation windows. Full MS scans were recorded at m/z 375–1455 with a resolution of 120,000 (at m/z 200), AGC target of 300%, and maximum injection time of 45 ms. MS2 scans were acquired at a resolution of 30,000 with normalized collision energies of 27, 30, and 32, AGC target of 1000%, automatic maximum injection time, and fixed first mass of m/z 200. Variable isolation windows (13.0, 16.0, 26.0, and 61.0 m/z) were applied with loop counts of 27, 13, 8, and 6, respectively.

#### 5.6.4. nanoLC-MS/MS analysis for phosphoproteomics

Phosphopeptides were analyzed on an Exploris 480 mass spectrometer (Thermo Scientific) coupled to a Vanquish Neo nano UPLC system with an EASY-Spray ion source. Samples were first trapped on an Acclaim PepMap 100 C18 trap column (75 µm × 2 cm, 3 µm, 100 Å, nanoViper) and separated on an EASY-Spray RSLC C18 analytical column (75 µm × 25 cm, 2 µm, 100 Å) at a flow rate of 300 nL/min and column temperature of 45 °C. Chromatographic separation was achieved using a nonlinear gradient with mobile phases consisting of (A) 0.1% formic acid and (B) 0.1% formic acid in 80% acetonitrile. Solvent B was increased from 5% to 25% over 110 min, then to 32% over 10 min, to 45% over 8 min, and finally to 95% in 2 min, which was maintained for 13 min. Data were acquired in data-dependent acquisition (DDA) mode using a top 15 method. Full MS1 scans were acquired at m/z 350–1750 with a resolution of 120,000 (at m/z 200), AGC target of 3 × 10^6, and maximum injection time of 45 ms. The 15 most intense precursor ions were selected for higher-energy collisional dissociation (HCD) with a normalized collision energy of 27. MS2 scans were recorded at a resolution of 45,000 with AGC target of 1 × 10^5, maximum injection time of 120 ms, and a dynamic exclusion of 30 s.

### 5.7. Raw data processing

#### 5.7.1. Metabolomics

GC–MS data were exported from ChromaTOF in NetCDF format and processed in MATLAB R2021a using custom scripts for baseline correction, chromatogram alignment, compression, and multivariate curve resolution. Metabolites were annotated based on retention indices and mass spectral matching against reference libraries such as Schauer et al^94^ and NIST MS Search Program v2.2 (National Institute of Standards and Technology, Gaithersburg, MD, USA), with ±5 RI units as the acceptance threshold. LC–MS data were processed in Agilent MassHunter Profinder (vB.10.0.2) using the Batch Targeted Feature Extraction workflow (20 ppm m/z tolerance; 0.1 min RT window). An in-house library of authentic standards and plant metabolites acquired under identical conditions was used for annotation, integrating MS, MS/MS, and retention time information.

Confidence levels for annotation of the metabolites were assigned according to established community systems^95–97^. Five levels were defined: Level 1 (confirmed identification), based on direct comparison with an authentic reference standard analyzed under identical conditions, requiring agreement in accurate mass, retention time, and usually MS/MS spectra; Level 1* (isomeric mixture), applied when standards confirmed a mixture of structural isomers without chromatographic or MS/MS separation; Level 2 (putative annotation/structure), based on spectral library matches and/or diagnostic MS/MS evidence but without a standard; Level 3 (tentative annotation/compound class), based on accurate mass, isotope distribution, and database search, enabling tentative structure or class assignment; Level 4 (molecular formula), features with accurate mass and molecular formula but insufficient evidence for structure; and Level 5 (unique feature), accurate mass only, without confirmed formula.

#### 5.7.2. Proteomics

MS files were analyzed using Spectronaut vs16.2 (Biognosys AG). A combined DDA-DIA spectral library was generated, containing 175364 precursors, 125493 peptides, and 9767 protein groups. DIA files were searched against this library using factory default settings with MS1-level quantification. Carbamidomethylation on cysteine was set as a fixed modification, while oxidation on methionine residues was considered as a variable modification. A maximum of two missed cleavages was allowed. False discovery rate (FDR) control was applied at 1% both the peptide and protein levels.

#### 5.7.3. Phosphoproteomics

Phosphopeptide identification and label-free quantification was performed by means of the Proteome Discoverer v2.5 (Thermo Scientific) using SEQUEST HT as the search engine and a human protein database download from UniProt on 2020-11-09 (42304 sequences with isoforms. For the search, trypsin was selected as the protease, two missed cleavages were allowed, the tolerance was fixed at 10 ppm for MS1 and 0.02 Da for MS2, carbamidomethyl-cysteine was set as a static modification, while methionine oxidation, phosphorylation on serine, threonine, and tyrosine, and protein N-terminal acetylation were selected as dynamic modifications. The ptmRS algorithm was used for scoring phosphorylation sites, considering site probability threshold >75. Peptides and corresponding proteins were identified with 1% of FDR.

### 5.8. Data preprocessing and statistical analysis

Data preprocessing and statistical analyses were conducted in R (version 4.4.2). First, analytes were filtered for a minimum of three valid values in at least one condition. After data filtering, each dataset was normalized using variance stabilizing normalization with the vsn R package^98^ (version 3.77.0). To account for technical variation across experimental runs, batch effects were corrected using the removeBatchEffect function from the limma package^99^ (version 3.58.0). Batch assignment and relevant covariates including sex, MS injection order, age, and postmortem brain interval (for PMBT samples) were included in the model to adjust expression values while preserving biological differences between experimental groups. Batch-corrected expression matrices were used for downstream unsupervised and multivariate analyses, including principal component analysis, Orthogonal Partial Least Squares Discriminant Analysis (OPLS-DA), and Weighted gene co-expression network analysis (WGCNA). Differential expression analyses were performed separately on vsn-normalized data using linear models with covariate adjustment as described below. Principal component analysis was conducted using the mixOmics package^100^ (version 6.26.0), for quality assessment.

Differential expression analysis was performed using linear models with empirical Bayes moderation as implemented in the limma package^99^ (version 3.58.0). Variance-stabilized normalized (vsn) data were used as input, with sex, age, and post-mortem brain interval (for PMBT samples), and mass spectrometry injection batch included as covariates. Proteins and phosphopeptides with a threshold of ± log2FC ≥ 0.5 and p-value < 0.05 were considered dysregulated.

To explore the features that separate each disease group from controls in the batch-corrected metabolomics dataset, orthogonal projections to latent structures discriminant analysis (predI = 1, orthoI = 1, crossvalI = 6) was performed using the ropls R package^101^ (version 1.34.0). The 75th percentile was used as the cutoff to select the top contributing features.

Weighted gene co-expression network analysis (WGCNA) was performed on normalized, batch-corrected matrices using the WGCNA R package^102^ (version 1.73) along with supporting packages such as tidyverse^103^ (version 2.0.0), magrittr (version 2.0.3), fastcluster^104^ (version 1.2.6), and dynamicTreeCut^105^ (version 1.63-1). The analysis was conducted with multi-threading enabled to optimize computational performance. A range of soft-thresholding powers (1–20) was evaluated using the pickSoftThreshold function to identify the lowest power at which scale-free topology was approximated (R² ≥ 0.90). A power of 9 was selected for network construction. Signed adjacency and topological overlap matrices (TOMs) were computed using bicorrelation as the similarity measure. Modules of co-expressed analytes were identified via hierarchical clustering and the dynamic tree-cutting method, with a minimum module size of 30 analytes and a merge cut height of 0.25. Module eigengenes were subsequently calculated for module-trait association analyses. To examine associations between module eigengenes and experimental traits, condition and grouping variables were incorporated into downstream statistical analyses. Non-parametric tests were employed to assess significance: Kruskal–Wallis tests were applied to detect overall differences across multiple conditions (p < 0.05). Where significant, Dunn’s post-hoc tests (p < 0.05) were performed for pairwise comparisons. Mann–Whitney U tests were used for binary group comparisons (p < 0.05). Multiple testing correction was performed using the Benjamini–Hochberg procedure to control the false discovery rate (FDR). Boxplots of module eigengene expression values across groups were used to illustrate significant associations, with significance levels indicated by standard thresholds (*p < 0.05, **p < 0.01, ***p < 0.001). All statistical analyses and visualizations were conducted in R using ggplot2^106^ (version 3.5.2) and ggpubr (version 0.6.1).

### 5.9. Kinome analysis

To investigate upstream kinase–substrate relationships, phosphosites identified in our dataset were annotated using publicly available databases. First, we searched the UniProt Knowledgebase to determine whether kinases had been previously reported for each phosphosite. We also applied the PhosphoSitePlus®^107^ (v6.8.1) resource to identify candidate kinases. Predicted kinase–substrate associations were prioritized by considering the balance of PhosphoSitePlus® scoring metrics, including the log2(score), score rank, site percentile, and percentile rank. These candidate kinase-substrate relationships were further evaluated by considering: 1) whether the candidate kinases were quantified in the proteomic dataset, and 2) the subcellular localization of both substrates and candidate kinases using information from UniProt^108^ and the Human Protein Atlas^109^. Predicted interactions were retained when the kinase and substrate were reported or predicted to localize to compatible subcellular compartments, ensuring biological plausibility.

### 5.10. Biological pathway analyses

Biological pathway enrichment analyses were performed separately for proteomic, phosphoproteomic, and metabolomic datasets. For proteins and phosphopeptides, pathway enrichment was carried out using the Functional Annotation Tool in DAVID Bioinformatics Resources (https://davidbioinformatics.nih.gov) and the STRING database (https://string-db.org). Analyses were performed against Reactome Gene Sets and KEGG Pathways, and significance was assessed using Fisher’s exact test. Pathways with nominal p < 0.05 were considered significantly enriched. For metabolomic data, enrichment analysis was conducted using MetaboAnalyst (https://www.metaboanalyst.ca) via the pathway analysis module. Pathways with nominal p < 0.05 were retained.

### 5.11. Concordance analysis between iPSC spheroids and postmortem brain tissue

Disease-associated effects were calculated as the median differences relative to control samples (ΔVSN = disease - control) within each dataset and disease group for iPSC spheroids and PMBT. Group median-difference values were used to increase robustness and reduce sensitivity to outliers. Comparative analyses between sample types were restricted to molecular features detected as dysregulated in at least one sample type. Global concordance of disease-associated molecular effects between sample types was assessed using directional concordance defined as sign concordance across shared features. To evaluate the robustness of the concordance metrics, non-parametric bootstrap resampling was performed at the molecular-feature level using bootstrap R (S-Plus). Shared features were resampled with replacement, and concordance metrics were recalculated across 5,000 iterations. Empirical 95% confidence intervals were derived from the resulting bootstrap distributions. Differences in disease-associated effects between sample types were quantified using absolute differences between iPSC spheroids and PMBT. Molecular features were subsequently classified according to both effect size and direction to define concordant and discordant molecular patterns.

## Supporting information

Velasquez Roybon Figure_S1

Velasquez Roybon Figure_S2

Velasquez Roybon Figure_S3

Velasquez Roybon Figure_S4

Velasquez Roybon Figure_S5

Velasquez Roybon Figure_S6

Velasquez Roybon Figure_S7

Velasquez Roybon Supporting_Figure_Legends

Velasquez Roybon Table_S1_Sample_information

Velasquez Roybon Table_S2_Cell type markes in spheroids

Velasquez Roybon Table_S3_Normalized protein intensities in spheroids

Velasquez Roybon Table_S4_Dysregulated proteins in spheroids

Velasquez Roybon Table_S5_Normalized metabolite intensities in spheroids

Velasquez Roybon Table_S6_Dysregulated metabolites in spheroids

Velasquez Roybon Table_S7_Normalized phosphopeptide intensities in spheroids

Velasquez Roybon Table_S8_Dysregulated phosphopeptides in spheroids

Velasquez Roybon Table_S9_Normalized proteins, metabolites and phosphopeptides intensities in PMBT

Velasquez Roybon Table_S10_Dysregulated proteins, metabolites and phosphopeptides in PMBT

## Resource availability

- **Lead contact**. Laurent Roybon
- **Material availability**. This study used iPSC lines that are subject to general data protection regulation. Brain samples were obtained from the Netherlands Bran Bank (https://www.brainbank.nl/).
- **Data availability**. All omics datasets are provided as supplementary material. Additionally, the mass spectrometry proteomics data have been deposited to the ProteomeXchange Consortium via the PRIDE^110^ partner repository with the dataset identifiers PXD074353 and PXD074573.

## Acknowledgments

We thank Dr. Stefano Goldwurm from the Parkinson Institute, ASST PINI-CTO, Milan, Italy (https://www.asst-pini-cto.it/), and the members of the Roybon laboratory at Lund University for their help with the generation of tools and reagents. We acknowledge the Cell Line and DNA Biobank From Patients Affected by Genetic Diseases, Istituto G. Gaslini, Genova, Italy, and the Parkinson Institute Biobank, both members of the Telethon Network of Genetic Biobanks funded by Telethon Italy (project no. GTB12001, http://biobanknetwork.telethon.it), for providing fibroblast samples. We are also thankful to the Netherlands Brain Bank for providing patient postmortem tissue samples. We also acknowledge the Swedish National Infrastructure for Biological Mass Spectrometry (BioMS) for providing access and support with mass spectrometry analyses. This work was supported by Swedish private foundations to L.R., including the Holger Crafoord Foundation, the Shaking Generation Foundation, the Petrus and Augusta Hedlunds Foundation, the Åke Wiberg Foundation, the Greta och Johan Kocks Foundation, donations for science, medicine, and technology at Fysiografen in Lund; the Swedish Research Council (Grants VR-2015-03684 and VR-2021-02284 to L.R.); Brainstem, Stem Cell Center of Excellence in Neurology, funded by the Innovation Fund Denmark, to L.R., and the Olav Thon Foundation in Norway, to L.R. We are also thankful to the Van Andel Institute for its support to L.R. and A.C.B.

## Author contributions

- **Conceptualization:** L.R., E.V.
- **Methodology:** L.R., M.R., E.V., A.J., E.N, Y.P, E.S, A.C.B., K.L. T.D.
- **Analysis:** L.R., M.R., E.V., A.G.T., A.J., E.N, A.C.B.
- **Investigation:** L.R., M.R., E.V.
- **Resources:** L.R., M.R.
- **Writing:** L.R., M.R., E.V., A.G.T.
- **Supervision:** L.R., M.R.
- **Funding acquisition:** L.R.

## Supplemental information

The supplementary information are available online.

