## Supplementary figures and images for "A convergent metabolic-kinase signaling axis links Parkinson’s disease and multiple system atrophy"

### Velasquez Roybon Figure_S1

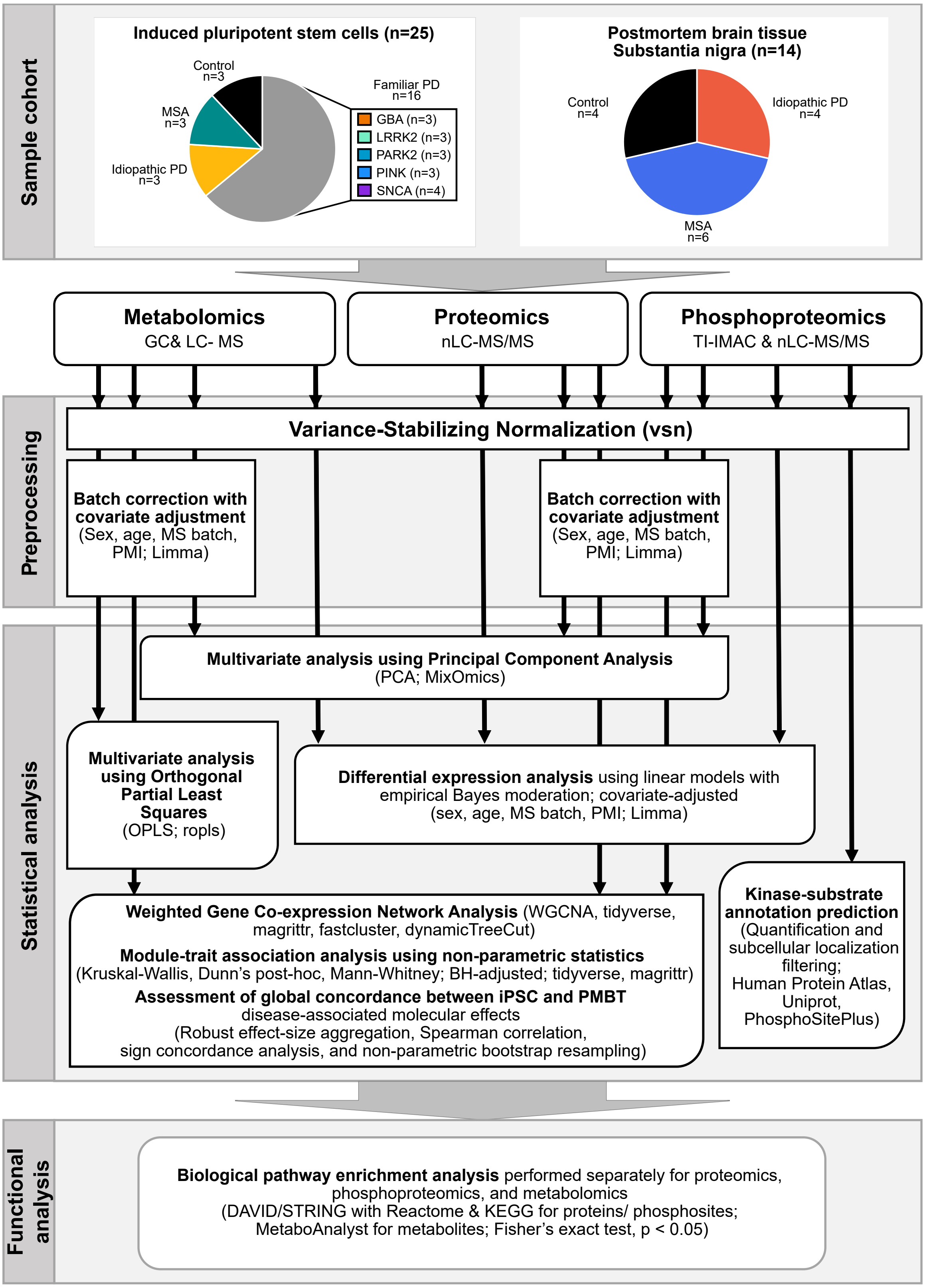

### Velasquez Roybon Figure_S2

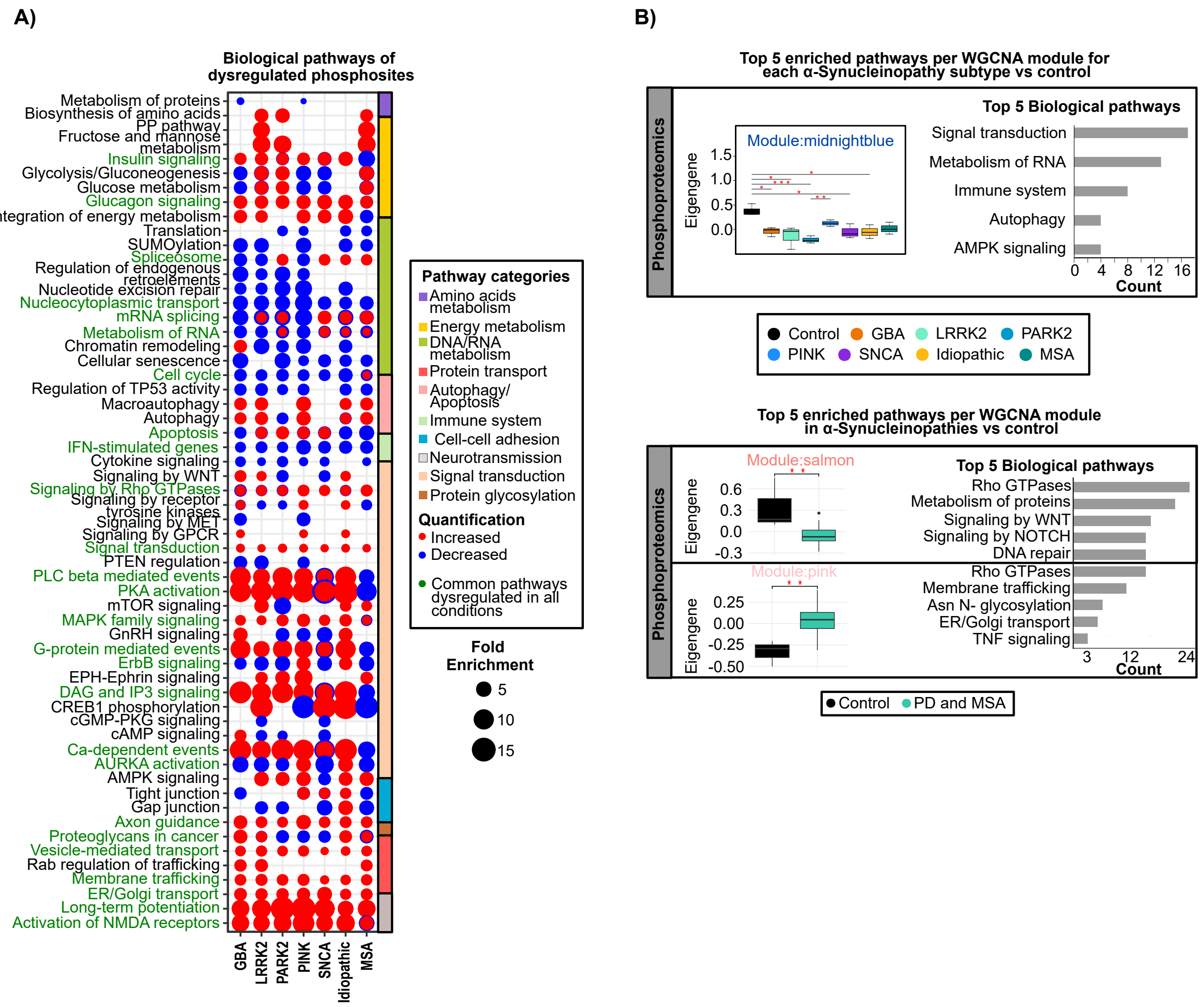

### Velasquez Roybon Figure_S3

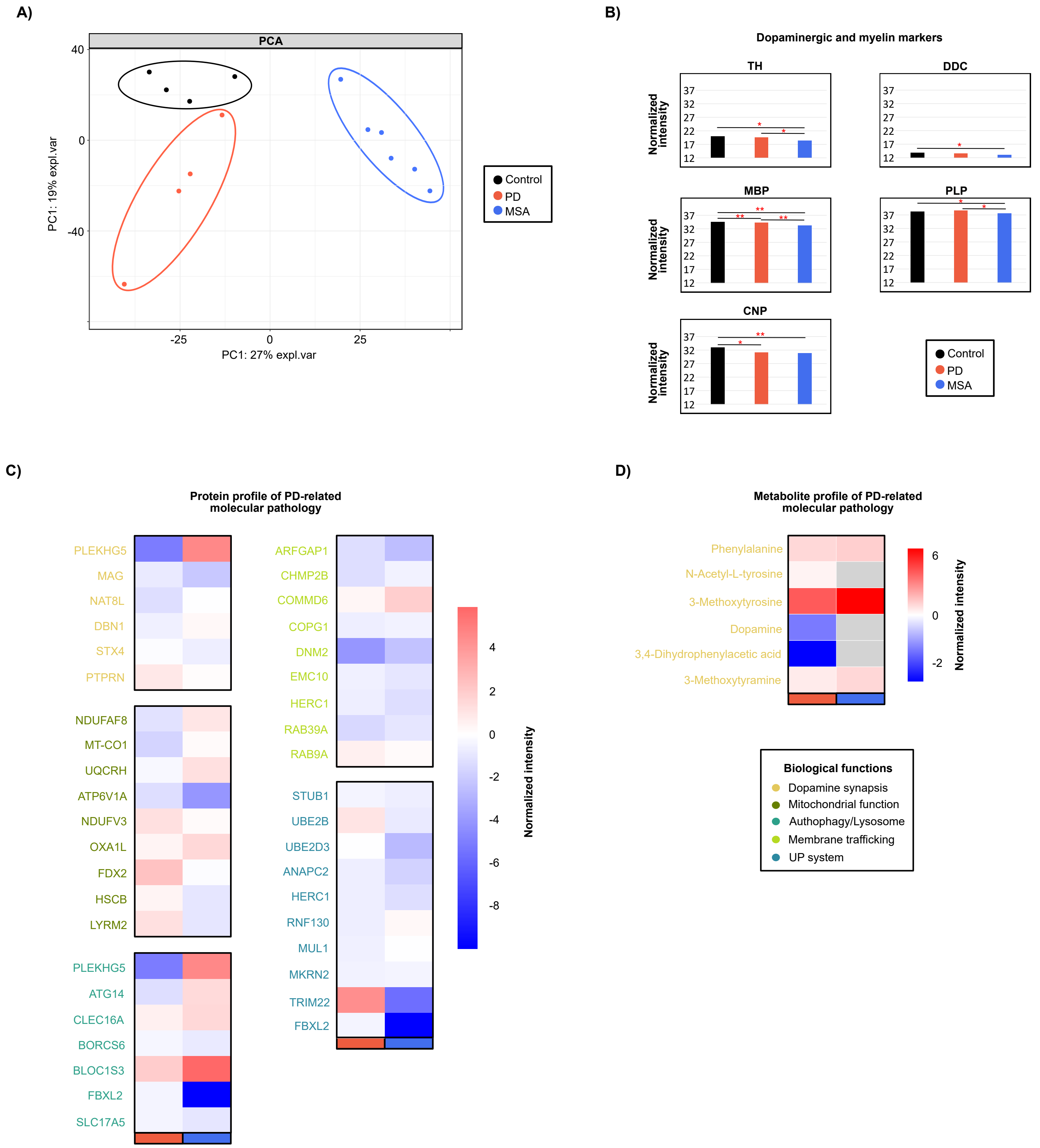

### Velasquez Roybon Figure_S4

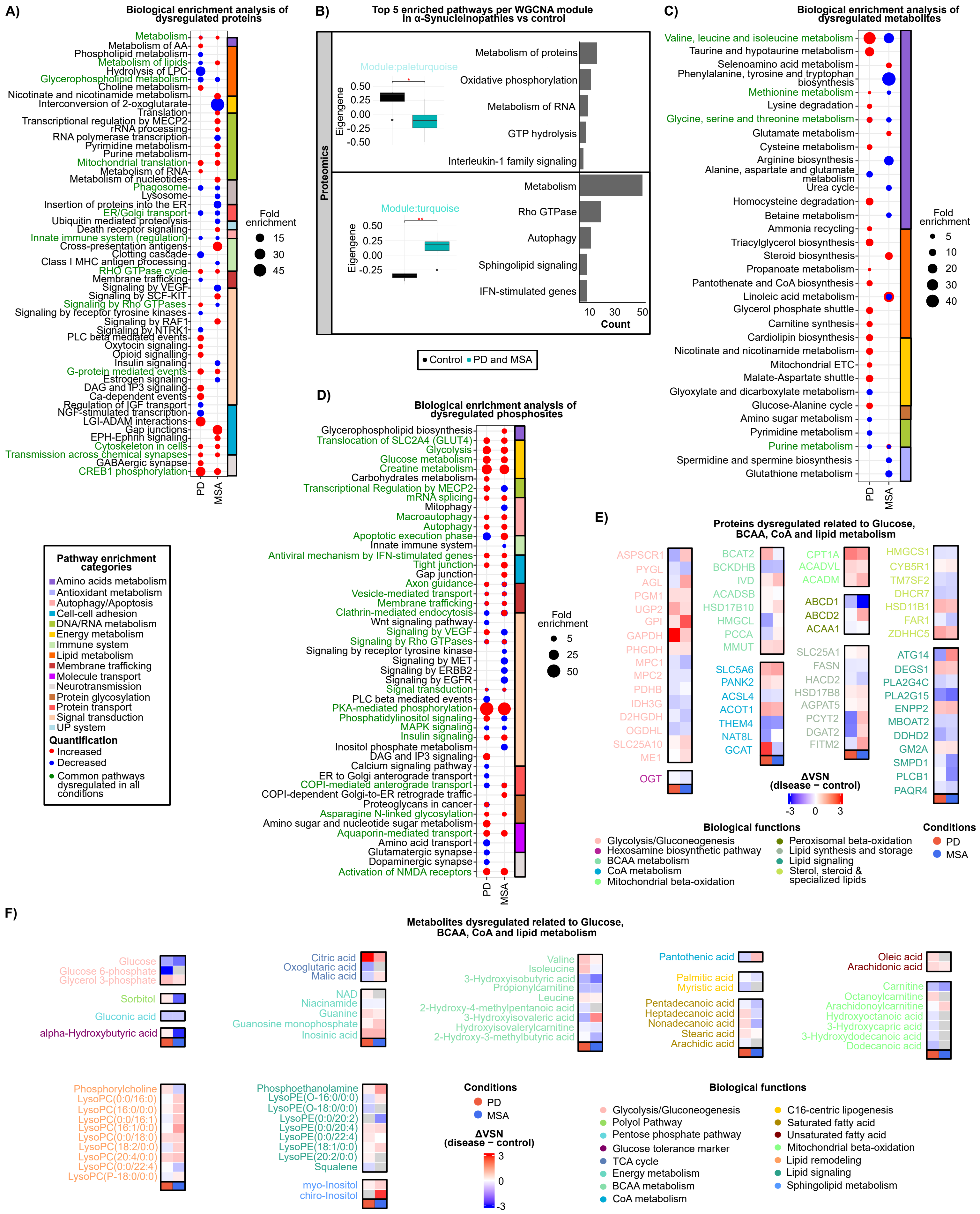

### Velasquez Roybon Figure_S5

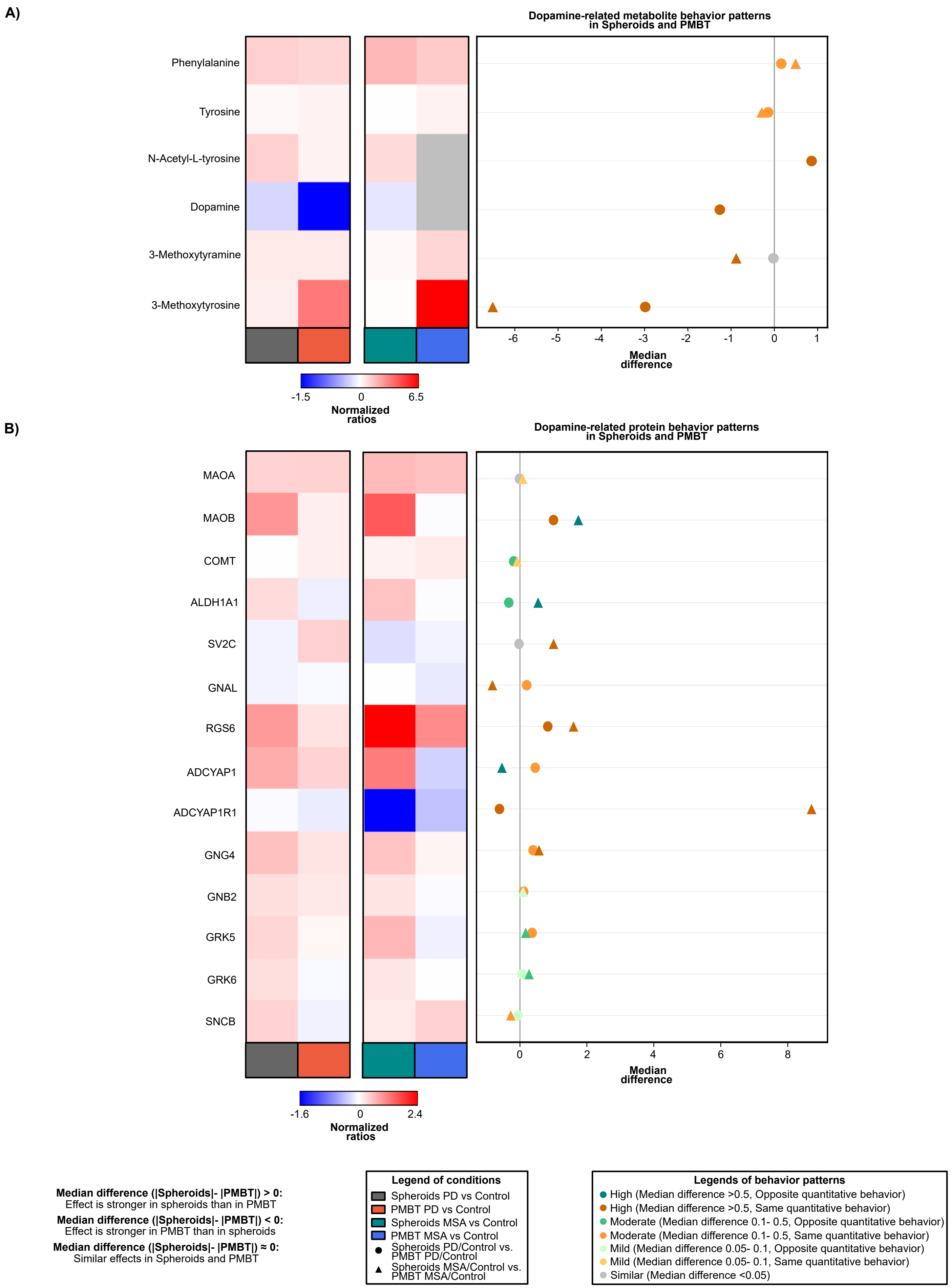

### Velasquez Roybon Figure_S6

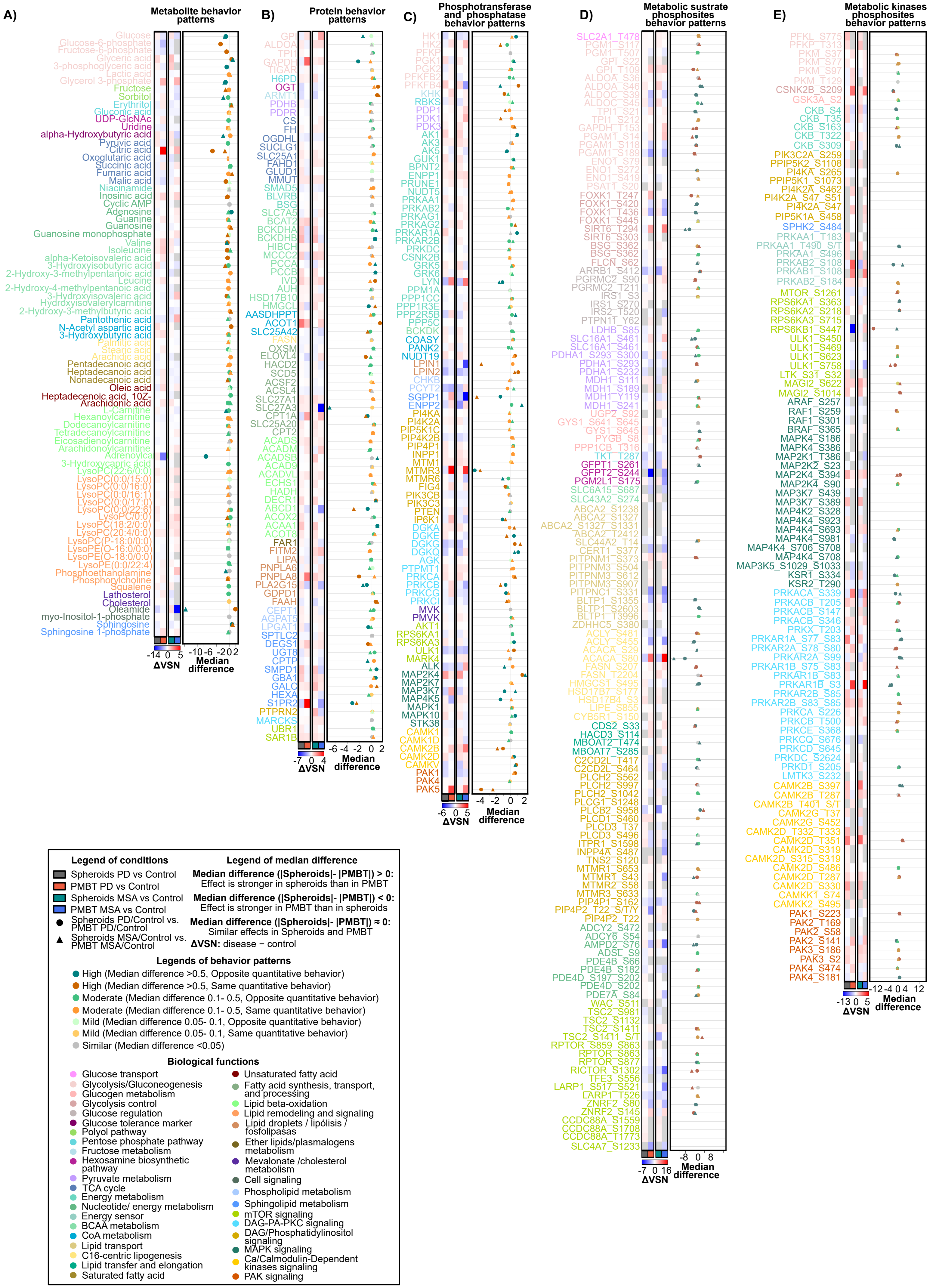

### Velasquez Roybon Figure_S7

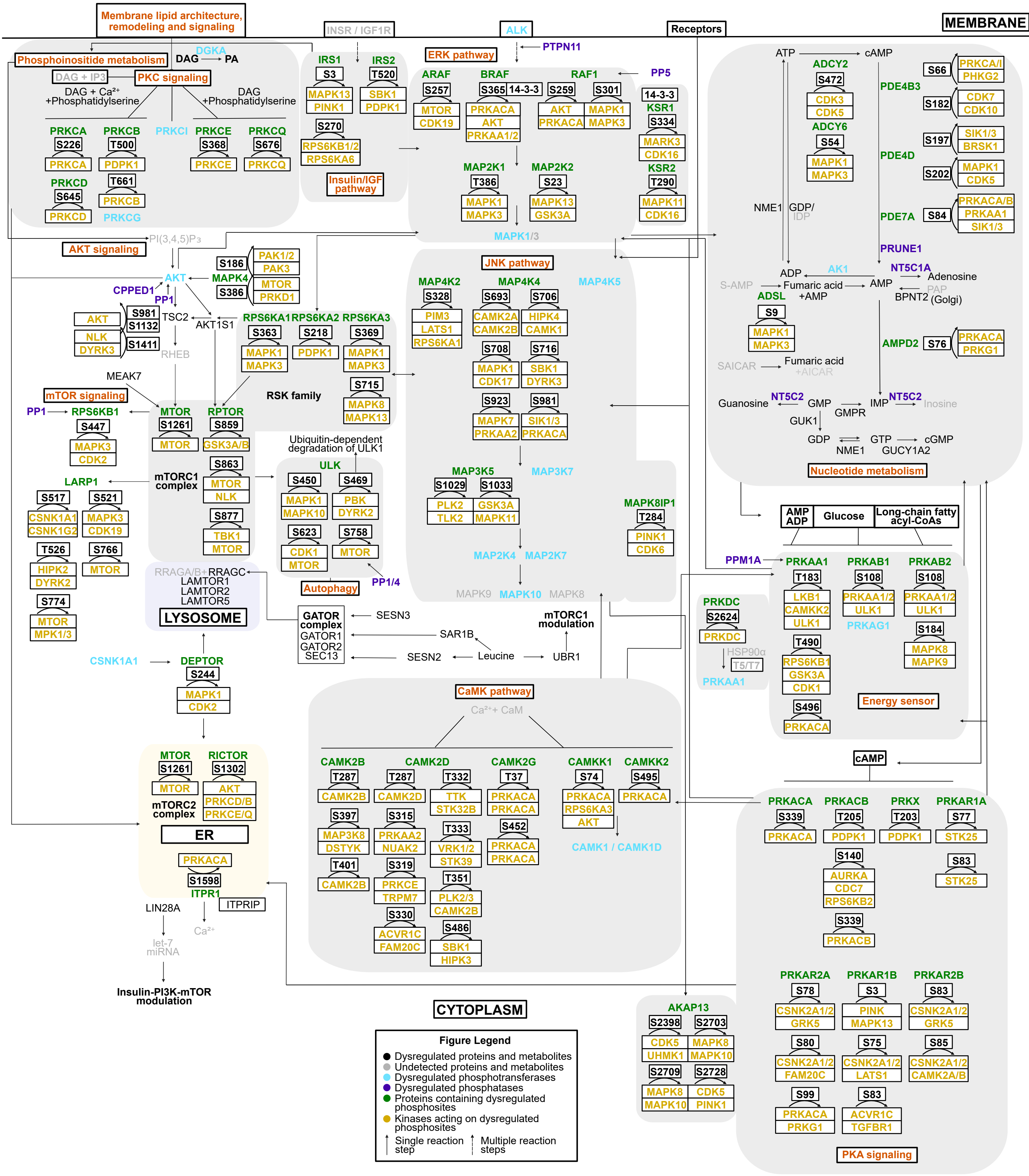
