## Supplementary material for "A convergent metabolic-kinase signaling axis links Parkinson’s disease and multiple system atrophy": Velasquez Roybon Supporting_Figure_Legends

**Supplementary Figure Legends**

**Figure S1. Overview of the analytical workflow used for sample preparation and downstream data analysis.**

**Figure S2. Phosphoproteomic pathway and module-level analysis in PD and MSA spheroids.** (A) Bubble plot showing pathway enrichment analysis of phosphopeptides identified as dysregulated in PD and MSA samples relative to controls. Dysregulated phosphopeptides were defined using robust empirical Bayes analysis (p < 0.05, |log2FC| ±0.5), and pathway enrichment was assessed using Fisher’s exact test (p < 0.05). Pathways highlighted in green indicate processes shared across all conditions. (B) Weighted gene co-expression network analysis (WGCNA) of phosphopeptide modules displaying significant differences among PD, MSA, and control groups (Kruskal-Wallis, p < 0.05; Mann–Whitney U test, p < 0.05). The five most significantly enriched biological pathways for each major module are shown (Fisher’s exact test, p < 0.05).

**Figure S3. Characterization of PMBT proteomic and metabolic alterations.** (A) Principal component analysis (PCA) of PMBT proteomes showing separation across the analyzed conditions. (B) Dopaminergic markers (TH, DDC) and myelin/oligodendroglial protein markers (MBP, PLP, CNP) identified as dysregulated using robust empirical Bayes analysis (p < 0.05, |log2FC| > ±0.5). (C) Annotated heatmap of PD-associated dysregulated proteins identified by robust empirical Bayes analysis (p < 0.05, |log2FC| > ±0.5). (D) Annotated heatmap of dopamine-related metabolites selected by OPLS analysis using a cutoff > 0.75.

**Figure S4. Multi-layer analysis of PMBT datasets.** (A) Bubble plot showing pathway enrichment analysis of proteins dysregulated in PD and MSA samples compared with controls. Dysregulated proteins were defined using robust empirical Bayes analysis (p < 0.05, |log2FC| ±0.5), and enriched pathways were identified using Fisher’s exact test (p < 0.05). Green-labeled pathways indicate shared dysregulation across all conditions. (B) Weighted gene co-expression network analysis (WGCNA) of protein modules showing significant differences across PD, MSA, and control groups (Mann–Whitney U test, p < 0.05). The top five enriched biological pathways for each major module are displayed (Fisher’s exact test, p < 0.05). (C–D) Bubble plots showing enrichment analysis of dysregulated (C) metabolites and (D) phosphopeptides in PD and MSA samples compared with controls (robust empirical Bayes, p < 0.05, |log2FC| ±0.5; Fisher’s exact test, p < 0.05). Pathways shown in green correspond to processes commonly altered across all conditions. (E–F) Dysregulated (E) proteins and (F) metabolites associated with glucose, BCAA, CoA, and lipid metabolism (robust empirical Bayes, p < 0.05, |log2FC| ±0.5; OPLS, cutoff > 0.75).

**Figure S5. Concordance analysis of dopaminergic metabolism between PD/MSA spheroids and substantia nigra postmortem brain tissue (PMBT).** Disease-associated effects were estimated as median differences relative to control samples within each dataset and disease group (VSN = disease − control). (A–B) Annotated heatmaps display the side-by-side quantification and behavioral patterns of dysregulated (A) metabolites and (B) proteins across spheroids and PMBT.

**Figure S6. Global concordance analysis between PD/MSA spheroids and substantia nigra postmortem brain tissue (PMBT)**. Disease-associated effects were calculated as median differences relative to control samples within each dataset and disease group (VSN = disease − control). (A–C) Annotated heatmaps show side-by-side quantification and behavioral patterns of dysregulated (A) metabolites, (B) proteins, and (C) phosphosites in PD and MSA spheroids and PMBT, emphasizing the principal metabolic processes affected across datasets.

**Figure S7. Integrated mapping of upstream kinase networks across spheroids and PMBT in PD and MSA.** All dysregulated metabolites, proteins, and phosphosites were integrated together with the top predicted kinases targeting these phosphosites. Kinases and molecular features associated with metabolic dysfunction in at least one condition were included to generate the integrated upstream kinase-network map.
